# A state with increased arousal threshold in *Araneus diadematus* (Araneidae) measured in the wild: new evidence for sleep in spiders

**DOI:** 10.1101/2023.10.19.563109

**Authors:** Daniela C. Rößler, Marie E. Herberstein

## Abstract

Sleep is a seemingly universal behavior across the animal kingdom, yet for the majority of species, experimental evidence thereof is still lacking. The recent report of REM sleep-like behavior in a jumping spider has highlighted the potential of spiders as a non-model organism to study invertebrate sleep. While behavioral evidence of potential sleep-states in spiders is strong, a crucial piece of evidence is so far lacking: a shift in arousability during sleep compared to awake states. Targeting a spider exquisitely suited for conducting experiments in the wild, we collected arousal threshold data for the diurnal orb-web spider *Araneus diadematus*. Our field experiments revealed significant differences in response latency between day- and night-times. Using a sound stimulus of 400 Hz with increasing amplitude that robustly triggers an anti-predatory response (raising of front legs), we tested both immobile and active spiders during the day and during the night. We found that spiders had a significantly longer response latency to the stimulus during nighttime immobility compared to immobile spiders during the day. There was no difference in the response latency between active spiders at night and active spiders during the day. Overall, our data demonstrate a shift in arousability during periods of night-time immobility in support of sleep in *A. diadematus*. Additionally, however, we found eight spiders that did not respond to the stimulus within the set time limit, most of which we encountered during daytime immobility hinting at additional sleep behavior during the day and a potential bimodal sleep pattern. Our study, along with recent work on jumping spider sleep-like behavior showcases the suitability of spiders for sleep research.

## Introduction

In recent years, the study of sleep in non-traditional animals has gained increasing interest from both sleep researchers as well as behavioral ecologists aimed at unraveling the origins and evolution of this intriguing behavior (Rattenborg & Ungurean, 2023). Although experimental evidence for sleep is lacking for the majority of species, there is a consensus that likely all animals engage in sleeping behavior. Sleep has been documented across several taxa from cnidarians, cephalopods, to arthropods (Hendricks et al., 2000; Iglesias et al., 2019; Nath et al., 2017; Shaw et al., 2000), reptiles, birds and mammals (Blumberg et al., 2020; Rattenborg & Ungurean, 2023; Shein-Idelson et al., 2016) with varying evidence for different sleep stages (REM sleep, Non-REM sleep, etc, Rattenborg & Ungurean, 2023).

Sleep is a behavioral state, set aside from wakefulness by an often-characteristic posture, immobility, reversibility of the state, homeostatic regulation as well as an increased arousal threshold (Campbell & Tobler, 1984; Piéron, 1913). Different sleep phases (e.g. REM sleep from non-REM sleep) can further be defined by changes in brain activities (e.g. Deboer, 2015).

While the best approach to experimentally establish sleep in an animal is to demonstrate multiple (or all) sleep criteria, showing differences in arousal thresholds between sleep and wake states is the gold standard. Other criteria such as posture or immobility can be considerably less reliable as, in some extreme cases, animals still move while sleeping (e.g. frigate birds sleep and fly at the same time by only engaging one brain half in the sleeping behavior, Rattenborg et al., 2016).

Sleep is particularly difficult to study in invertebrates due to their often-small size as well as different neural organization. Apart from model organisms such as *Drosophila*, comparatively few studies have been conducted on sleep in other invertebrates. These include cnidaria (Nath et al., 2017), molluscs (Vorster et al., 2014), nematodes (Trojanowski & Raizen, 2016), various cephalopods (Iglesias et al., 2019; Medeiros et al., 2021), bees (Kaiser & Steiner-Kaiser, 1983), crayfish (Osorio-Palacios et al., 2021) and scorpions (Tobler & Stalder, 1988) (list is not exhaustive). While sleep has been established in all of these animals, only cephalopods have so far been shown to demonstrate mammalian-like sleep architecture with distinct sleep phases resembling those of mammals including humans (Pophale et al., 2023). Sleep in spiders has recently been indicated but has not been experimentally established.

In a recent report, Rößler et al. (2022) documented nocturnal behavioral phases in jumping spiders reminiscent of REM sleep in mammals and other vertebrates. The phases included myoclonic twitching, apparent muscle atonia (demonstrated by curling of the legs likely due to a drop in hydraulic pressure) as well as retinal movements. The behaviors occurred periodically at robust lengths and intervals over the course of the night, when the spiders were immobile and adopted a characteristic resting position (Rößler et al., 2021). We have since recorded similar behaviors in numerous spider lineages (Geiger et al., *in prep*) including in orb-web spiders (e.g. *Araneus diadematus*). Similar to the original observation on jumping spiders (Rößler et al., 2022), we recorded the same behavioral markers constituting potential REM sleep-like phases in *A. diadematus* including leg curling (**Video S1**), twitching of abdomen, legs, pedipalps and spinnerets (**Video S2**). These conspicuous but short phases commenced around dusk (**Fig. 1**), while the spider resided in the hub of its web.

**Figure 1.**
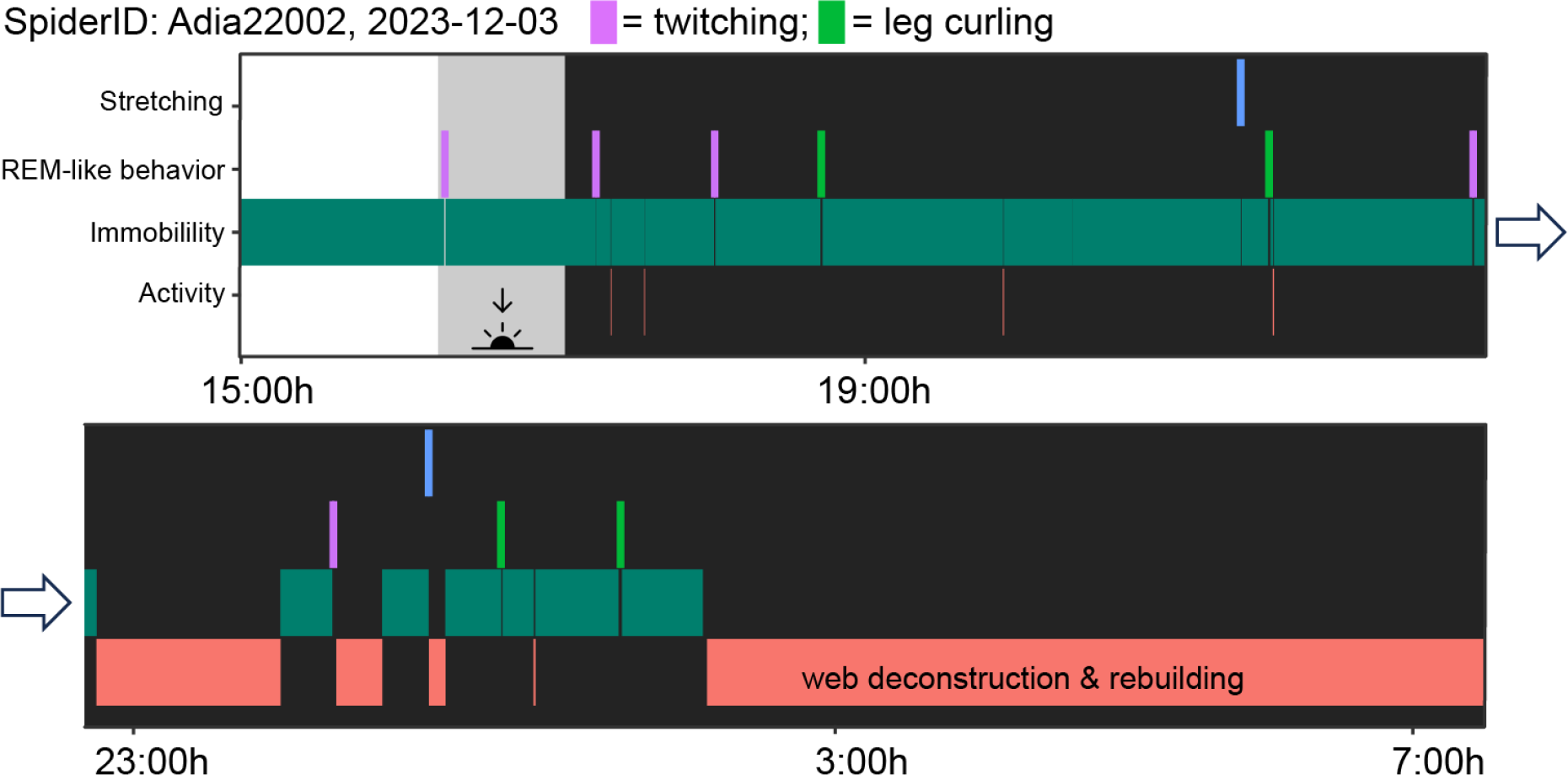
Ethogram of a penultimate female *Araneus diadematus* filmed over a 16h period starting in afternoon until the next morning in the laboratory (sunset at the time of recording was at 16:30h, grey background indicates twilight). The spider shows the longest immobility periods at the onset of the night together with several bouts of REM sleep-like behaviors (twitching = bouts of irregular twitching of pedipalps and spinnerets; leg curling = bouts with inward curling of multiple legs and irregular twitching). Prolonged activity starts around 3:00h with the deconstruction and rebuilding of the web. Bars showing twitching, leg curling and stretching have been widened for better visibility (Mean duration of REM sleep-like bouts = 44.99 s; SD = 19.21 s).

Additionally, we observed *A. diadematus* in the field and sometimes encountered individuals with noticeably different body postures such as a drooping abdomen (**Fig. 2B,C, Video S3**) or one or both pedipalps hanging visibly low (**Fig. 2D**). Upon touch these spiders would often not react, which hinted at the presence of sleep states with an increased arousal threshold. Based on these observations, our aim was to test experimentally whether *A. diadematus* show differences in arousal thresholds, in support of sleep.

**Figure 2.**
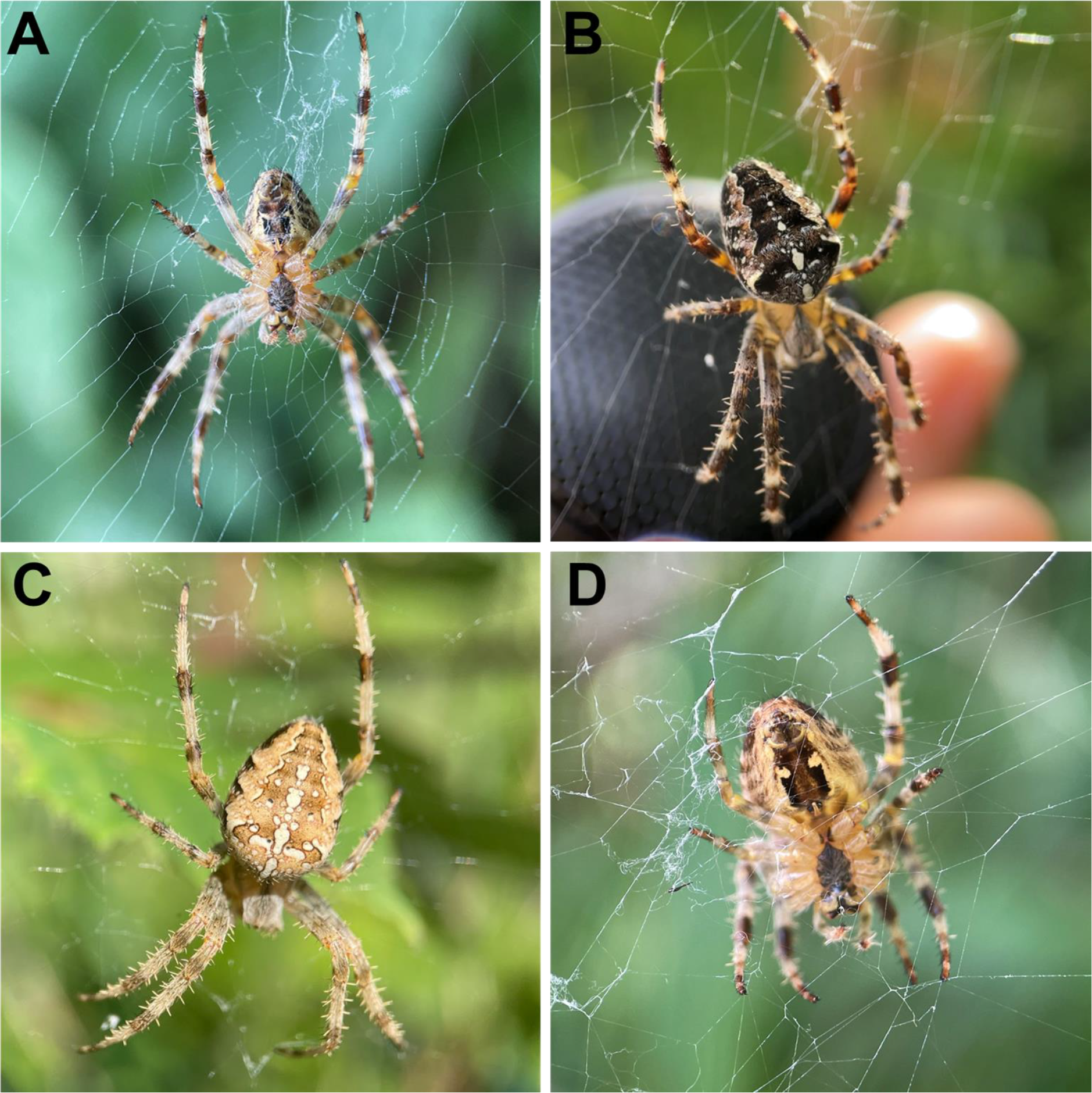
Examples of body postures of *Araneus diadematus* and response times encountered during the study. **A**: Spider A03 in an alert position with the spinnerets close (or touching) the web, legs symmetrically positioned. Response latency (immobile, day) = 2.1 s. **B**: Spider A47 with a visibly drooped abdomen. Bluetooth speaker in the background. **C**: Spider A12 with a visibly drooped abdomen and asymmetrical leg position. Spiders in B and C did not react towards the first stimulus presentation. **D**: Spider A04 with one pedipalp hanging visibly lower. Response latency (immobile, day) was 3.3 s. Drooping abdomen, asymmetrical leg position or lowered pedipalps could indicate relaxation of muscle tone. Photo credit: Daniela C. Rößler.

In doing so, we enthusiastically adopted the well-argued need for sleep studies in the wild (Loftus et al., 2022; Rattenborg et al., 2017; Reinhardt, 2020). Field sleep research is important to capture natural sleep duration and patterns, which are often inflated in simple and less demanding laboratory conditions (e.g. easily available food and limited interactions with other individuals or species; see Rattenborg et al. 2017). Drawbacks of field research include a lack of control over ambient conditions, as well as difficulties in tracking individuals across multiple hours or days.

A related issue is the choice of arousal stimuli. Various stimuli are mostly applied in the laboratory and while convenient to use, their ecological relevance is often not established. In our study, we opted for a stimulus that would generate a stereotypical anti-predator response, without signs of habituation. In sum, the high field density of *A. diadematus*, the abundance and ease of access to the spiders at the hub of the web, their stationary nature as well as their robust response to the test stimulus are an ideal combination for field experimentation on sleep.

Nevertheless, there are challenging aspects to the biology of *A. diadematus* requiring careful formulation of predictions for arousal threshold experiments. As a sit-and-wait predator, orb-web spiders do not have clearly distinguishable active and inactive phases, unlike jumping spiders (e.g. Rößler et al., 2021) or other non-traditional study systems (e.g. sharks, Kelly et al. 2020), because they spend the majority of their time immobile in the hub of their web. While the spiders are clearly in an active wakeful state, for example during feeding, grooming or web-building, we believe that immobility can entail both wakeful (i.e. vigilant), quiescence (ready to capture prey) as well as sleep-like states. Based on our preliminary observations of spiders engaging in leg-curling and twitching phases after sunset, we expect sleep-associated immobility to take place more frequently during the night and predict these to be characterized by an increased arousal threshold.

*Assumption 1*: immobility in the web may be associated with a sleep-like state.

*Assumption 2*: as the spiders’ main activity period is during the day, sleep-like states are more likely (but not exclusively) to occur during the night.

*Assumption 3:* response latency to a stimulus is an indicator of arousal threshold (e.g. Vorster et al. 2014): longer response latency indicates a higher arousal threshold, shorter response latency indicates a lower arousal threshold.

*Essential comparison*: to control for environmental differences between day and night such as temperature and visual input, or overall motivational state, the focus of comparing responses to a stimulus is between active and immobile states within each time period (day and night)

### Predictions

1. If spiders engage in sleeping behavior during the night, we predict that **during the night**, immobile spiders have a longer response latency to a stimulus than active spiders
2. If spiders are vigilant during the day, we predict that **during the day**, immobile spiders do not differ in response latency to a stimulus from active spiders
3. We predict that immobile spiders at night will have a higher response latency to a stimulus than immobile spiders during the day.
4. We predict no difference in the response latency to a stimulus between active spiders during the day and during the night.

To our knowledge, this is the first experimental field study explicitly testing arousal thresholds in an invertebrate.

## Methods

### Field site and study species

Field observations and experiments were conducted in August and September 2023 near the Schmugglerbucht, Lake Constance, Konstanz, Germany (47.665027, 9.199751). *Araneus diadematus* built their webs in shrub vegetation that lined the walking path along the lakefront. Prior to the beginning of the study, we systematically observed spider activity throughout different times of the day as well as at night. Our initial observations confirmed that during that time the majority of spiders resided at the hub of the web throughout the day and the night, deconstructing and rebuilding their webs in the early hours of the morning (∼4:00h). Based on these observations we consider that the predominant activity period for *A. diadematus* is during the day, when most of its foraging takes place while the web is fresh. Although *A*.

*diadematus* can also actively capture prey and feed during night time, the then often highly damaged webs due to daytime activity and the timing of web reconstruction in early morning hours accord with a diurnally-adapted optimal activity (Peters, 1970; Ramousse & Davis, 1976). We thus decided to conduct arousal tests during the day (between ∼9:00h and ∼17:00h) and late night (between 21:00 and ∼24:00h). We exclusively tested penultimate and adult females in our study. Adult males of this species are considerably smaller than adult females, and do not build webs. Adult and penultimate males can be easily distinguished from females.

### Stimulus design

To test for arousal thresholds, we used a biologically meaningful stimulus (a sound) that produced a robust anti-predatory response–spiders consistently raised their front legs (**Video S4**). This ‘rebuffing’ behavior is well known in orb-web spiders (Tolbert, 1975)and has been interpreted as the spiders pushing the threatening stimulus away with their front legs. This behavior can be induced using tuning forks at various frequencies that elicit species specific responses (see Davies & Hesselberg 2022 for a review). Our preliminary observations confirmed that the response is robustly triggered irrespective of whether it is presented dorsally or ventrally (see also Frings & Frings, 1966). After a number of pre-trials, we found that a sine wave of 400 Hz with increasing amplitude for 10 seconds robustly elicited a swift rebuffing behavior in *A. diadematus*. This frequency is within the range of wingbeat frequency bees and wasps produce (i.e., 100-580 Hz, Parmezan et al., 2021). Thus, a delayed response, or lack of response can carry significant fitness costs for the spider if approached by a predatory wasp. The sound was controlled via an iPhone 12 mini (Apple Inc., USA) by using the *Audio Function Generator* app (https://ee-toolkit.com/audio-function-generator/) which has an in-built amplitude-sweep function (set to 10 s and at a system volume of 50), administered via a mini Bluetooth speaker (Eageroo) held 4-5 cm from the body of the spider at the hub. Actual sound output from the bluetooth speaker (frequency and amplitude) was measured with an iPhone 12 mini using the app *Decibel X: dB Sound Level Meter* (SkyPaw Co. Ltd) with a response time of 0.2 s (dB graph, **Supplementary Data S1**). We consider the total stimulus intensity to be a product of stimulus strength (dB) and stimulus duration (**Fig. 3B**). We then measured the response latency to this stimulus (until the raising of the front legs) to the nearest millisecond by taking screenshots of the time display on the audio function generator app on the iPhone.

**Figure 3.**
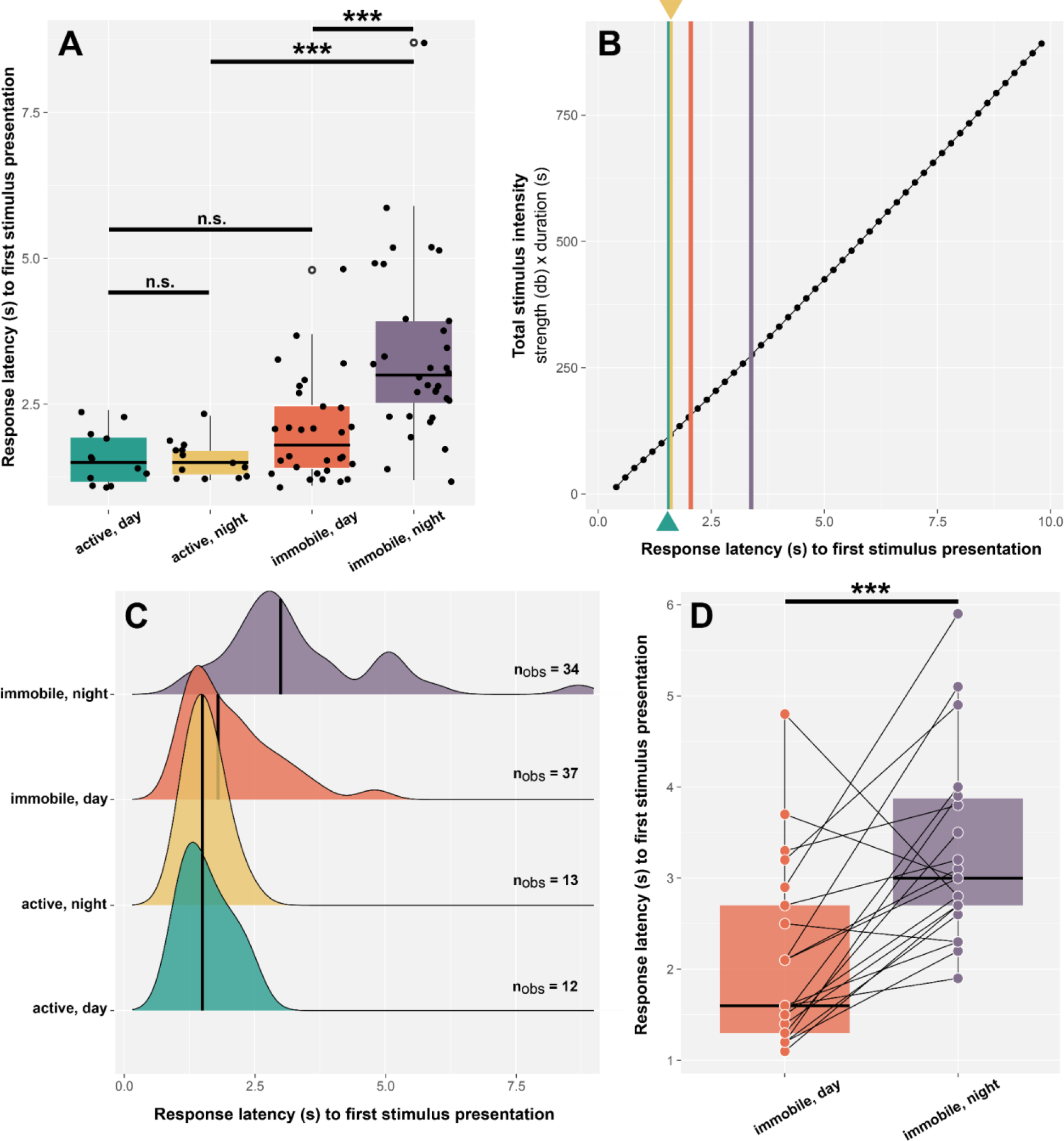
Response latency during different behaviors and daytimes. **A**: Boxplots showing response latency towards the first stimulus presentation depending on behavior and daytime. Black horizontal lines representing the median, lower and upper bound of the boxes show 25th and 75th percentiles with whiskers representing ±1.5 interquartile range. White circles represent outliers. **B**: Mean response latency plotted on the total stimulus intensity (stimulus strength (dB) x duration (s)), illustrating the arousal thresholds based on behavior and daytime. **C**: Smoothed density plots showing distribution of response latency. Black lines show medians. **D**: Boxplots showing response latency to first stimulus presentation only for retested individuals (20 spiders with two data points). Black lines connecting individual’s data points.

### Field experiment

Our experimental design was to test the response of individuals in active and immobile states (while in the web) during the day and during the night in the field. We located individuals and marked their website with flagging tape and assigned an ID. A subset of individuals was first tested during the day and the night second. The other half of individuals were first tested during the night and then during the day. There was a period of at least 6h between tests. At the time of testing, we noted if the spider was active (feeding, cleaning, web repair) or immobile (no visible movement of any body part for several minutes prior to testing). While other studies testing arousal thresholds (e.g. sharks: Kelly et al., 2021) use the term ‘inactive’, we prefer to use the term immobile, as orb-web spiders, being sit-and-wait predators (Herberstein, 2011) are generally immobile when at the hub. Thus, immobility can entail both wakeful and quiet states.

Due to the unpredictable nature of animals in the field, we were not able to retrieve all individuals and test them in all conditions (active/immobile; day/night). A total of 23 individual spiders were retrieved and retested during immobile day and night conditions, three of which did not respond towards the first stimulus presentation in one of the tests (**Fig. 3D**). For a subset of 28 spiders, after the stimulus presentation, we measured their size (body length) by gently removing each spider from the hub and placing it in a large (14.5 cm) petri dish and photographing it against a scale. We then gently returned each spider to the hub of their web. Images were then used to measure body length (spinnerets to anterior rim of the prosoma) using the software ImageJ (Schneider et al., 2012). We located spiders at night using a dimmed headlamp to reduce light exposure. We did not observe any visible reaction to the dim light during our study.

Unexpectedly, we came across some individuals who did not respond to the stimulus within the 10 second presentation period (**Fig. 2B,C**). We considered whether these individuals were unresponsive because they a) were in a phase of deep sleep, b) disregarded the stimulus be it due to inability to perceive the stimulus or because they deemed it non-threatening. Consequently, we used a blade of grass to engage the spider, by gently touching its body until the spider visibly moved. Once the spider resettled in the hub, we waited for 20 s and presented the stimulus again recording the response latency as described before.

### Statistical Analysis

Statistical analyses were carried out in R 4.2.1 (R Core Team, 2023). We used GLMMs using the package *glmmTMB* (Magnusson et al., 2017). We first checked for potential effects of spider size. To test which factors had significant effects on the dependent variables, we then applied an analysis of deviance to the resulting models using the package *car* (Fox & Weisberg, 2018). Spider ID was always included as a random factor to account for repeated measures. Post hoc analyses with Bonferroni correction were conducted using the package *emmeans* (Lenth et al., 2018). Model fit was confirmed using the package *DHARMa* (Hartig, 2017). We modeled the response latency towards the stimulus as influenced by daytime and behavior using a Gamma error distribution with a log link. All plots were generated using the package *ggplot2* (Wickham, 2016). All data and the full R script containing the data analysis are available from the supplementary material (**Supplementary Information S1, Supplementary Data S1-S3**).

## Results

In total, we tested 96 stimulus presentations with 67 individuals. We found no effect of body size (GLMM analysis of deviance, χ^2^ = 0.093, p = 0.759, n_obs_ = 49, n_sub_ = 28), or of test order (day or night first) on the response latency towards the first stimulus presentation (GLMM analysis of deviance, χ^2^ = 1.497, p = 0.221, n_obs_ = 40, n_sub_ = 20). Overall, behavior (active/immobile) during different time periods (day/night) significantly predicted response latency towards the first stimulus presentation (GLMM analysis of deviance, χ^2^ = 62.22, p < 0.0001, n_obs_ = 87, n_sub_ = 61).

### Prediction 1

#### during the night, immobile spiders have a longer response latency than active spiders

Spiders immobile during the night had significant longer response latency than spiders active during the night (Post-hoc, Bonferroni corrected, z.ratio = -6.023, SE = 0.058, p < 0.0001).

### Prediction 2

#### during the day, immobile spiders do not differ in response latency from active spiders

Spiders active during the day and immobile during the day did not differ in response latency (Post-hoc, Bonferroni corrected, z.ratio = -1.665, SE = 0.1, p = 0.575).

### Prediction 3

#### immobile spiders at night will have a longer response latency than immobile spiders during the day

Immobile spiders during the night had a longer response latency than immobile spiders during the day (GLMM, post-hoc, Bonferroni corrected, z.ratio = -6.034, SE = 0.049, p < 0.0001).

### Prediction 4

#### no difference in response latency between active spiders during the day or the night

Spiders that were active during the day or night did not differ in response latency (GLMM, post-hoc, Bonferroni corrected, z.ratio = 0.181, SE = 0.142, p = 1.00). Of the 96 tests conducted, nine spiders did not respond within 10 seconds to the stimulus at its first presentation. To test if these non-responders did not perceive the test stimulus as threatening or were unable to perceive it, we re-tested them after gentle activation. Only one of the nine spiders did not react to the second stimulus presentation after activation and was therefore excluded from all analyses. The response latencies of the other eight spiders were within the range of diurnal response latency of immobile spiders (**Table 1**). Among the eight spiders that did not initially respond, one was displaying leg curling behavior (A21).

**Table 1.** Response latency of eight *Araneus diadematus* individuals that did not react to the initial stimulus presentation, after gentle activation and repeated exposure to the stimulus.

| Spider ID | behavior, daytime | response latency (s) after activation |
| --- | --- | --- |
| A02 | immobile, **day** | 3.8 |
| A12 | immobile, **day** | 1.2 |
| A47 | immobile, **day** | 1.6 |
| A40 | immobile, **day** | 1.2 |
| A43 | immobile, **day** | 2.7 |
| A48 | immobile, **day** | 1.8 |
| A64 | immobile, **night** | 2.9 |
| A21 | immobile, **night** | 1.9 |

Non-responders made up 8.42% of all trials and exclusively occurred during immobile behavioral states. They constituted 5.56% of immobile night trials and 13.95% of immobile day trials. Including a maximum response time of 10s for non-responders in the response latency did not change the overall picture of the data analysis: Overall, behavior (active/immobile) during different time periods (day/night) still significantly predicted response latency towards the first stimulus presentation (GLMM analysis of deviance, χ^2^ = 32.008, p < 0.0001, n_obs_ = 95, n_sub_ = 67). Post-hoc analysis still revealed a significant difference between immobile spiders during the day and immobile spiders during the night: GLMM, post-hoc, Bonferroni corrected, z.ratio = -2.958, SE = 0.082, p = 0.0186)

## Discussion

In our study we found that the interaction between activity state and time of day had a strong effect on response latency towards a sound stimulus in *Araneus diadematus* orb-web spiders. We found no effects of body size or test order on the response latency towards the stimulus. Immobile spiders at night took significantly longer to react to the stimulus than active spiders at night as well as compared to immobile spiders during the day, however, we found a higher number of initially unresponsive spiders during daytime immobility, suggestive of the presence of daytime sleep associated with a considerable increase of arousal threshold. Overall, during the day, spiders responded similarly, regardless of whether they were immobile or active. Moreover, active spiders at night responded similarly to active spiders during the day.

As we set out in the introduction, we predicted that immobile spiders at night are more likely to engage in sleep-like behavior and should show greater response latency to the stimulus, a prediction supported by our data. There are of course a number of non-sleep state explanations why spiders during the night might respond differently than during the day. Ambient temperature is typically lower at night; thus, spiders might respond slower than during the day. Similarly, predation pressure might also be lower at night (see Rypstra 1984), visual input is much reduced at night, or spider motivation during the night is just different for other unknown reasons.

Our field experiment was designed to minimize alternative explanations as we also measured the response latency of nighttime active spiders as a control. If lower temperatures or reduced predation pressure at nighttime triggers a general greater response latency, then we should see no difference between active and immobile spiders at night and a difference between day active and night active spiders. Our data do not support these alternative explanations as active spiders at night responded with almost the same latency than active spiders during the day (a peak of around 1.5 s) while immobile spiders at night took more than twice as long to respond (3.2 s). Our results are similar to the arousal thresholds in *Drosophila* that also compared thresholds of flies during the day and during the night in the absence of ambient light (see van Alphen et al. 2013). However, there may be unknown interactions between nighttime and vigilance that affect arousal thresholds, for example hunger or mating states, which would be interesting to explicitly test in future work.

One of the assumptions we set out is that response latency is an indicator of arousal threshold. In the sleep literature, arousal thresholds are tested by either response latency towards a static stimulus (e.g. Vorster et al., 2014) or by increasing stimulus strength until a response is detected (e.g. Kelly et al., 2021). The stimulus we tested combines these two approaches by measuring the response latency to an increasing stimulus intensity (Figure 3B). Based on previous studies, we are confident that the response latency we measured is an appropriate proxy for arousal threshold. Considering our assumptions, experimental design and unanimous support for our predictions together, we believe we have the strongest evidence to date that a spider engages in sleep behavior.

Immobile spiders during the day responded as quickly to the stimulus as active spiders during the day, suggesting that they were not in a sleep, but in a quiescent state. However, among immobile spiders during the day we see a longer tail in the frequency distribution with a small number of individuals showing delayed response times that match the frequency distribution pattern of immobile spiders at night suggesting that these spiders also enter a sleep-like state during the day. This is further supported by our finding that some spiders did not respond at all within the set time frame of 10s. That “no responses” exclusively occurred during immobility states further supports that immobility can be associated with both awake as well as asleep phases. Six out of the eight non-responding spiders were tested during daytime immobility. Thus, it is highly likely that diurnal *A. diadematus* engage in a bimodal sleep pattern similar to *Drosophila*, which shows both daytime and nighttime sleep, where daytime sleep is shorter and arousal thresholds are lower compared to nighttime sleep (Huber et al., 2004; Ishimoto et al., 2012). Interestingly, our small sample of non-responding spiders during the day would indicate the opposite to be the case for these spiders, with daytime sleep entailing a higher arousal threshold. Possible explanations could include the difference of sensory environment between daytime and nighttime or the need to enter a deeper sleep stage during the day. Another interesting question here is how spiders optimally allocate sleep and awake behavior during the day and the night relative to other fitness generating behaviors such as foraging and predator avoidance.

The picture that emerges from our study here and previous work (Rößler et al., 2021, 2022) is that a sleep state is present in spiders. To date, the presence of different sleep stages has been established in mammals, birds, reptiles and cephalopods (Pophale et al., 2023; Rattenborg & Ungurean, 2023; Shein-Idelson et al., 2016). To further confirm sleep stages does require electrophysiological experimentation that is notoriously difficult in spiders due to the hydrostatic pressure present in the carapace that holds the central nervous system. However, some exciting progress has been made in this field holding potential for future electrophysiological research in spiders (Menda et al., 2014; Shamble et al., 2016).

What is highly interesting is that REM sleep-like behaviors (leg curling, twitches) in A. *diadematus* in the laboratory have so far exclusively been observed in the dark or at twilight (**Fig. 1**). Considering the large number of eminent researchers over the last century who have spent hundreds of hours observing *A. diadematus* detailing its web-building, prey-capture and mating behavior, the only reasonable explanation for why a conspicuous behavior such as leg curling has not been described in detail, is that it is indeed an exclusively nocturnal behavior. From a predator risk perspective, uncontrollably twitching and leg curling is likely to increase conspicuousness of an otherwise well camouflaged spider. However, it is plausible that predation pressure for *A. diadematus* is lower at night in the absence of diurnal visually hunting predators such as wasps and birds (see Wise, 1993). Of note is that sleep-like behaviors such as leg curling and twitching in *A. diadematus* are near identical (though shorter, Geiger et al. *in prep*) to those reported in jumping spiders (Rößler et al., 2022), even though the two spider lineages are separated by at least 200 million years (Magalhaes et al., 2020).

As research of sleep in spiders is only in its infancy, we are of course cautious in our interpretation and enthusiastic to follow up our arousal threshold experiments with additional observations and experiments (on several spider lineages) that further establish a sleep state in spiders both in the laboratory as well as in the field. These include testing homeostatic regulation confirming that spiders experience increased sleep pressure after deprivation, (Campbell & Tobler, 1984), establishing behavioral markers and especially behavioral proxies (i.e., postures) of sleep, and explicitly testing arousal during different sleep phases (e.g. REM sleep-like bouts) in more controlled environments.

Beyond the establishment of sleep in spiders, we also argue that spiders are particularly excellent models where sleep behavior can be integrated into wakeful behavior to explain the well-established variation in aspects of foraging behavior (e.g. web size and architecture, Herberstein & Heiling, 1999, 2001) and mating behavior (e.g. sexual cannibalism and paternity biases e.g. Elgar et al., 2000; Heiling & Herberstein, 2000; Herberstein et al., 2005, 2014; Schneider et al., 2000). Moreover, the diverse ecological niches spiders occupy (e.g. nocturnal/diurnal; active hunters/sit-and-wait predators; vison/vibratory focused) could facilitate a strong contribution by spiders towards linking ecology with sleep function (Lesku et al., 2012; Lesku & Rattenborg, 2022).

## Acknowledgements

We would like to thank Barrett Klein for valuable feedback on earlier versions of our manuscript.

## Inclusion and diversity statement

The authors greatly value equity, diversity and inclusion (EDI) in science. This study was conducted under consideration of EDI best practice. The authors represent different career stages (postdoc and professor). One or more of the authors self-identifies as a member of the LGBTQ+ community. We actively worked to promote gender balance (of first author) in our reference list, which, however, we did not fully reach.

## References

van Alphen, B., Yap, M. H. W., Kirszenblat, L., Kottler, B., & van Swinderen, B. (2013). A dynamic deep sleep stage in Drosophila. The Journal of Neuroscience, 33(16), 6917–6927.

Blumberg, M. S., Lesku, J. A., Libourel, P.-A., Schmidt, M. H., & Rattenborg, N. C. (2020). What Is REM Sleep? Current Biology: CB, 30(1), R38–R49.

Campbell, S. S., & Tobler, I. (1984). Animal sleep: a review of sleep duration across phylogeny. Neuroscience and Biobehavioral Reviews, 8(3), 269–300.

Davies, M. S., & Hesselberg, T. (2022). The use of tuning forks for studying behavioural responses in orb web spiders. Insects, 13(4).

Deboer, T. (2015). Behavioral and electrophysiological correlates of sleep and sleep homeostasis. Current Topics in Behavioral Neurosciences, 25, 1–24.

Elgar, M. A., Schneider, J. M., & Herberstein, M. E. (2000). Female control of paternity in the sexually cannibalistic spider Argiope keyserlingi. Proceedings. Biological Sciences / The Royal Society, 267(1460), 2439–2443.

Fox, J., & Weisberg, S. (2018). An R Companion to Applied Regression. SAGE Publications.

Frings, H., & Frings, M. (1966). Reactions of Orb-Weaving Spiders (Argiopidae) to Airborne Sounds. Ecology, 47(4), 578–588.

Hartig, F. (2017). DHARMa: residual diagnostics for hierarchical (multi-level/mixed) regression models. R package. Vienna, Austria: CRAN. https://CRAN.R-project.org/package=DHARMa.

Heiling, A. M., & Herberstein, M. E. (2000). Interpretations of orb-web variability: A review of past and current ideas. https://www.european-arachnology.org/esa/wp-content/uploads/2015/08/097-106_Heiling.pdf

Hendricks, J. C., Finn, S. M., Panckeri, K. A., Chavkin, J., Williams, J. A., Sehgal, A., & Pack, A. I. (2000). Rest in Drosophila is a sleep-like state. Neuron, 25(1), 129–138.

Herberstein, M. E. (2011). Spider Behaviour: Flexibility and Versatility. Cambridge University Press.

Herberstein, M. E., Gaskett, A. C., Schneider, J. M., Vella, N. G. F., & Elgar, M. A. (2005). Limits to male copulation frequency: Sexual cannibalism and sterility in St Andrew’s cross spiders (Araneae, Araneidae). Ethology: Formerly Zeitschrift Fur Tierpsychologie, 111(11), 1050–1061.

Herberstein, M. E., & Heiling, A. M. (1999). Asymmetry in spider orb webs: a result of physical constraints? Animal Behaviour, 58(6), 1241–1246.

Herberstein, M. E., & Heiling, A. M. (2001). Positioning at the hub: does it matter on which side of the web orb-web spiders sit? Journal of Zoology, 255(2), 157–163.

Herberstein, M. E., Wignall, A. E., Hebets, E. A., & Schneider, J. M. (2014). Dangerous mating systems: signal complexity, signal content and neural capacity in spiders. Neuroscience and Biobehavioral Reviews, 46 Pt 4, 509–518.

Huber, R., Hill, S. L., Holladay, C., Biesiadecki, M., Tononi, G., & Cirelli, C. (2004). Sleep homeostasis in Drosophila melanogaster. Sleep, 27(4), 628–639.

Iglesias, T. L., Boal, J. G., Frank, M. G., Zeil, J., & Hanlon, R. T. (2019). Cyclic nature of the REM sleep-like state in the cuttlefish Sepia officinalis. The Journal of Experimental Biology, 222(Pt 1), jeb174862.

Ishimoto, H., Lark, A., & Kitamoto, T. (2012). Factors that differentially affect daytime and nighttime sleep in Drosophila melanogaster. Frontiers in Neurology, 3, 24.

Kaiser, W., & Steiner-Kaiser, J. (1983). Neuronal correlates of sleep, wakefulness and arousal in a diurnal insect. Nature, 301(5902), 707–709.

Kelly, M. L., Spreitzenbarth, S., Kerr, C. C., Hemmi, J. M., Lesku, J. A., Radford, C. A., & Collin, S. P. (2021). Behavioural sleep in two species of buccal pumping sharks (Heterodontus portusjacksoni and Cephaloscyllium isabellum). Journal of Sleep Research, 30(3), e13139.

Lenth, R., Singmann, H., Love, J., Buerkner, P., & Herve, M. (2018). Emmeans: Estimated marginal means. AKA Least-Squares Means.

Lesku, J. A., & Rattenborg, N. C. (2022). The missing cost of ecological sleep loss. Sleep Advances : A Journal of the Sleep Research Society, 3(1), zpac036.

Lesku, J. A., Rattenborg, N. C., Valcu, M., Vyssotski, A. L., Kuhn, S., Kuemmeth, F., Heidrich, W., & Kempenaers, B. (2012). Adaptive sleep loss in polygynous pectoral sandpipers. Science, 337(6102), 1654–1658.

Loftus, J. C., Harel, R., Núñez, C. L., & Crofoot, M. C. (2022). Ecological and social pressures interfere with homeostatic sleep regulation in the wild. eLife, 11.

Magalhaes, I. L. F., Azevedo, G. H. F., Michalik, P., & Ramírez, M. J. (2020). The fossil record of spiders revisited: implications for calibrating trees and evidence for a major faunal turnover since the Mesozoic. Biological Reviews of the Cambridge Philosophical Society, 95(1), 184–217.

Magnusson, A., Skaug, H., Nielsen, A., Berg, C., Kristensen, K., Maechler, M., van Bentham, K., Bolker, B., Brooks, M., & Brooks, M. M. (2017). Package “glmmTMB.” R Package Version 0. 2. 0. http://cran.uni-muenster.de/web/packages/glmmTMB/glmmTMB.pdf

Medeiros, S. L. de S., Paiva, M. M. M. de, Lopes, P. H., Blanco, W., Lima, F. D. de, Oliveira, J. B. C. de, Medeiros, I. G., Sequerra, E. B., de Souza, S., Leite, T. S., & Ribeiro, S. (2021). Cyclic alternation of quiet and active sleep states in the octopus. iScience, 24(4), 102223.

Menda, G., Shamble, P. S., Nitzany, E. I., Golden, J. R., & Hoy, R. R. (2014). Visual perception in the brain of a jumping spider. Current Biology: CB, 24(21), 2580–2585.

Nath, R. D., Bedbrook, C. N., Abrams, M. J., Basinger, T., Bois, J. S., Prober, D. A., Sternberg, P. W., Gradinaru, V., & Goentoro, L. (2017). The jellyfish Cassiopea exhibits a sleep-like state. Current Biology: CB, 27(19), 2984–2990.e3.

Osorio-Palacios, M., Montiel-Trejo, L., Oliver-Domínguez, I., Hernández-Falcón, J., & Mendoza-Ángeles, K. (2021). Sleep phases in crayfish: relationship between brain electrical activity and autonomic variables. Frontiers in Neuroscience, 15, 694924.

Parmezan, A. R. S., Souza, V. M. A., Žliobaitė, I., & Batista, G. E. A. P. A. (2021). Changes in the wing-beat frequency of bees and wasps depending on environmental conditions: a study with optical sensors. Apidologie, 52(4), 731–748.

Peters, P. J. (1970). Orb web construction: Interaction of spider (Araneus diadematus) and thread configuration. Animal Behaviour, 18, 478–484.

Piéron, H. (1913). Le problème physiologique du sommeil. Masson.

Pophale, A., Shimizu, K., Mano, T., Iglesias, T. L., Martin, K., Hiroi, M., Asada, K., Andaluz, P. G., Van Dinh, T. T., Meshulam, L., & Reiter, S. (2023). Wake-like skin patterning and neural activity during octopus sleep. Nature.

Ramousse, R., & Davis, F., 3rd. (1976). Web-building time in a spider: preliminary applications of ultrasonic detection. Physiology & Behavior, 17(6), 997–1000.

Rattenborg, N. C., de la Iglesia, H. O., Kempenaers, B., Lesku, J. A., Meerlo, P., & Scriba, M. F. (2017). Sleep research goes wild: new methods and approaches to investigate the ecology, evolution and functions of sleep. Philosophical Transactions of the Royal Society of London. Series B, Biological Sciences, 372(1734).

Rattenborg, N. C., & Ungurean, G. (2023). The evolution and diversification of sleep. Trends in Ecology & Evolution, 38(2), 156–170.

Rattenborg, N. C., Voirin, B., Cruz, S. M., Tisdale, R., Dell’Omo, G., Lipp, H.-P., Wikelski, M., & Vyssotski, A. L. (2016). Evidence that birds sleep in mid-flight. Nature Communications, 7, 12468.

Reinhardt, K. D. (2020). Wild primate sleep: understanding sleep in an ecological context. Current Opinion in Physiology, 15, 238–244.

Rößler, D. C., De Agrò, M., Biundo, E., & Shamble, P. S. (2021). Hanging by a thread: unusual nocturnal resting behaviour in a jumping spider. Frontiers in Zoology, 18(1), 23.

Rößler, D. C., Kim, K., & De Agrò, M. (2022). Regularly occurring bouts of retinal movements suggest an REM sleep–like state in jumping spiders. Proceedings of the. https://www.pnas.org/doi/abs/10.1073/pnas.2204754119

Rypstra, A. L. (1984). A relative measure of predation on web-spiders in temperate and tropical forests. Oikos, 43(2), 129–132.

Schneider, C. A., Rasband, W. S., & Eliceiri, K. W. (2012). NIH Image to ImageJ: 25 years of image analysis. Nature Methods, 9(7), 671–675.

Schneider, J. M., Herberstein, M. E., De Crespigny, F. C., Ramamurthy, S., & Elgar, M. A. (2000). Sperm competition and small size advantage for males of the golden orb-web spider Nephila edulis. Journal of Evolutionary Biology, 13(6), 939–946.

Shamble, P. S., Menda, G., Golden, J. R., Nitzany, E. I., Walden, K., Beatus, T., Elias, D. O., Cohen, I., Miles, R. N., & Hoy, R. R. (2016). Airborne Acoustic Perception by a Jumping Spider. Current Biology: CB, 26(21), 2913–2920.

Shaw, P. J., Cirelli, C., Greenspan, R. J., & Tononi, G. (2000). Correlates of sleep and waking in Drosophila melanogaster. Science, 287(5459), 1834–1837.

Shein-Idelson, M., Ondracek, J. M., Liaw, H.-P., Reiter, S., & Laurent, G. (2016). Slow waves, sharp waves, ripples, and REM in sleeping dragons. Science, 352(6285), 590–595.

Tobler, I., & Stalder, J. (1988). Rest in the scorpion — a sleep-like state? Journal of Comparative Physiology A, 163(2), 227–235.

Tolbert, W. W. (1975). Predator avoidance behaviors and web defensive structures in the orb weavers Argiope aurantia and Argiope trifasciata (Araneae, Araneidae). Psyche; a Journal of Entomology, 82(1), 29–52.

Trojanowski, N. F., & Raizen, D. M. (2016). Call it Worm Sleep. Trends in Neurosciences, 39(2), 54–62.

Vorster, A. P. A., Krishnan, H. C., Cirelli, C., & Lyons, L. C. (2014). Characterization of sleep in Aplysia californica. Sleep, 37(9), 1453–1463.

Wickham, H. (2016). ggplot2: Elegant Graphics for Data Analysis. Springer, Cham.

Wise, D. H. (1993). Spiders in Ecological Webs. Cambridge University Press.

