## Supplementary Information S1 for "A state with increased arousal threshold in *Araneus diadematus* (Araneidae) measured in the wild: new evidence for sleep in spiders"

2023-09-14

#### Load packages

```
library(ggplot2) #plots
library(readr)#write
library(DHARMa)#model fit
library(car) #anova
library(emmeans) #estimated marginal means
library(glmmTMB) #glmms
library(psych) #stats
library(reshape2) #make density plots
library(ggridges) #density plots
```

#### Set a colorblind friendly color palette

```
##Color Palette for plots
cbbPalette <- c("#E69F00", "#56B4E9", "#009E73", "#CC79A7")

# To use for fills, add scale_fill_manual(values=cbbPalette)
# To use for line and point colors, add scale_colour_manual(values=cbbPalette)
```

#### Load data

```
db <- read.csv("F:/AraneusArousal/SupplementaryData/SupplementaryData S1.csv") # dB measures/stimulus i
a <- read.csv("F:/AraneusArousal/SupplementaryData/SupplementaryData S2.csv") #main data set
rcap <- read.csv("F:/AraneusArousal/SupplementaryData/SupplementaryData S3.csv") #recapture data
```

#### Describe data

```
describeBy(a, a$behavior_daytime)
```

```
##
##  Descriptive statistics by group
## group: active_day
##                  vars  n    mean    sd  median trimmed   mad     min     max
## date*               1 12    3.00  0.00    3.00    3.00  0.00    3.00    3.00
## temp                2 12   24.00  0.00   24.00   24.00  0.00   24.00   24.00
## location*           3 12    1.00  0.00    1.00    1.00  0.00    1.00    1.00
## time                4 12 1530.17 59.81 1532.50 1529.60 74.13 1448.00 1618.00
## daytime*            5 12    1.00  0.00    1.00    1.00  0.00    1.00    1.00
## subject*            6 12   43.17  5.17   43.00   43.10  6.67   36.00   51.00
## behavior*           7 12    1.00  0.00    1.00    1.00  0.00    1.00    1.00
## behavior_daytime*   8 12    1.00  0.00    1.00    1.00  0.00    1.00    1.00
## first               9 12    0.00  0.00    0.00    0.00  0.00    0.00    0.00
## size_cm            10  1   11.73    NA   11.73   11.73  0.00   11.73   11.73
## reaction           11 12    1.00  0.00    1.00    1.00  0.00    1.00    1.00
## responselatency1   12 12    1.58  0.47    1.50    1.55  0.59    1.10    2.40
## reaction2          13  0     NaN    NA      NA     NaN    NA     Inf    -Inf
## responselatency2   14  0     NaN    NA      NA     NaN    NA     Inf    -Inf
## pot_sleep          15  0     NaN    NA      NA     NaN    NA     Inf    -Inf
## comments*          16 12    2.83  0.58    3.00    3.00  0.00    1.00    3.00
##                  range  skew kurtosis    se
## date*              0.0   NaN      NaN  0.00
## temp               0.0   NaN      NaN  0.00
## location*          0.0   NaN      NaN  0.00
## time             170.0  0.06    -1.48 17.27
## daytime*           0.0   NaN      NaN  0.00
## subject*          15.0  0.12    -1.56  1.49
## behavior*          0.0   NaN      NaN  0.00
## behavior_daytime*  0.0   NaN      NaN  0.00
## first              0.0   NaN      NaN  0.00
## size_cm            0.0    NA       NA    NA
## reaction           0.0   NaN      NaN  0.00
## responselatency1   1.3  0.48    -1.37  0.14
## reaction2         -Inf    NA       NA    NA
## responselatency2  -Inf    NA       NA    NA
## pot_sleep         -Inf    NA       NA    NA
## comments*          2.0 -2.65     5.48  0.17
## ------------------------------------------------------------
## group: active_night
##                  vars  n    mean    sd median trimmed   mad    min    max
## date*               1 13    3.00  0.00    3.0    3.00  0.00    3.0    3.0
## temp                2 13   19.00  0.00   19.0   19.00  0.00   19.0   19.0
## location*           3 13    1.00  0.00    1.0    1.00  0.00    1.0    1.0
## time                4 13 2136.00 24.59 2131.0 2131.73 14.83 2114.0 2205.0
## daytime*            5 13    2.00  0.00    2.0    2.00  0.00    2.0    2.0
## subject*            6 13   52.54  8.27   55.0   53.09  4.45   37.0   62.0
## behavior*           7 13    1.00  0.00    1.0    1.00  0.00    1.0    1.0
## behavior_daytime*   8 13    2.00  0.00    2.0    2.00  0.00    2.0    2.0
## first               9 13    0.69  0.48    1.0    0.73  0.00    0.0    1.0
## size_cm            10  0     NaN    NA     NA     NaN    NA    Inf   -Inf
## reaction           11 13    1.00  0.00    1.0    1.00  0.00    1.0    1.0
## responselatency1   12 13    1.55  0.33    1.5    1.52  0.30    1.2    2.3
## reaction2          13  0     NaN    NA     NA     NaN    NA    Inf   -Inf
## responselatency2   14  0     NaN    NA     NA     NaN    NA    Inf   -Inf
```

```
## pot_sleep           15  0     NaN    NA     NA     NaN    NA    Inf   -Inf
## comments*           16 13    3.31  0.75    3.0    3.18  0.00    3.0    5.0
##                   range  skew kurtosis   se
## date*               0.0   NaN      NaN 0.00
## temp                0.0   NaN      NaN 0.00
## location*           0.0   NaN      NaN 0.00
## time               91.0  1.61     1.95 6.82
## daytime*            0.0   NaN      NaN 0.00
## subject*           25.0 -0.84    -0.87 2.29
## behavior*           0.0   NaN      NaN 0.00
## behavior_daytime*   0.0   NaN      NaN 0.00
## first               1.0 -0.74    -1.56 0.13
## size_cm            -Inf    NA       NA   NA
## reaction            0.0   NaN      NaN 0.00
## responselatency1    1.1  0.72    -0.39 0.09
## reaction2          -Inf    NA       NA   NA
## responselatency2   -Inf    NA       NA   NA
## pot_sleep          -Inf    NA       NA   NA
## comments*           2.0  1.70     0.99 0.21
## ------------------------------------------------------------
## group: immobile_day
##                   vars  n    mean     sd  median trimmed   mad    min    max
## date*                1 36    2.28   1.21    3.00    2.23  1.48   1.00    4.0
## temp                 2 36   23.17   1.89   24.00   23.50  0.00  19.00   24.0
## location*            3 36    1.00   0.00    1.00    1.00  0.00   1.00    1.0
## time                 4 36 1480.61 248.67 1579.00 1513.43 72.65 927.00 1707.0
## daytime*             5 36    1.00   0.00    1.00    1.00  0.00   1.00    1.0
## subject*             6 36   24.03  18.38   20.00   22.03 17.79   1.00   67.0
## behavior*            7 36    2.00   0.00    2.00    2.00  0.00   2.00    2.0
## behavior_daytime*    8 36    3.00   0.00    3.00    3.00  0.00   3.00    3.0
## first                9 36    0.36   0.49    0.00    0.33  0.00   0.00    1.0
## size_cm             10 29   11.80   1.67   11.61   11.70  1.41   8.71   16.4
## reaction            11 36    0.83   0.38    1.00    0.90  0.00   0.00    1.0
## responselatency1    12 30    2.05   0.88    1.80    1.92  0.74   1.10    4.8
## reaction2           13  6    1.00   0.00    1.00    1.00  0.00   1.00    1.0
## responselatency2    14  6    2.05   1.02    1.70    2.05  0.74   1.20    3.8
## pot_sleep           15  4    1.00   0.00    1.00    1.00  0.00   1.00    1.0
## comments*           16 36    1.17   0.85    1.00    1.00  0.00   1.00    6.0
##                    range  skew kurtosis    se
## date*               3.00  0.04    -1.71  0.20
## temp                5.00 -1.71     0.97  0.31
## location*           0.00   NaN      NaN  0.00
## time              780.00 -1.44     0.46 41.44
## daytime*            0.00   NaN      NaN  0.00
## subject*           66.00  0.84    -0.17  3.06
## behavior*           0.00   NaN      NaN  0.00
## behavior_daytime*   0.00   NaN      NaN  0.00
## first               1.00  0.55    -1.74  0.08
## size_cm             7.69  0.77     0.70  0.31
## reaction            1.00 -1.71     0.97  0.06
## responselatency1    3.70  1.22     1.13  0.16
## reaction2           0.00   NaN      NaN  0.00
## responselatency2    2.60  0.68    -1.36  0.42
## pot_sleep           0.00   NaN      NaN  0.00
```

```
## comments*           5.00  5.21    26.74  0.14
## ------------------------------------------------------------
## group: immobile_night
##                   vars  n    mean    sd  median trimmed   mad     min    max
## date*                1 34    2.15  0.36    2.00    2.07  0.00    2.00    3.0
## temp                 2 34   18.15  0.36   18.00   18.07  0.00   18.00   19.0
## location*            3 34    1.00  0.00    1.00    1.00  0.00    1.00    1.0
## time                 4 34 2252.56 48.96 2242.00 2253.82 39.29 2146.00 2333.0
## daytime*             5 34    2.00  0.00    2.00    2.00  0.00    2.00    2.0
## subject*             6 34   24.24 17.55   22.50   22.36 14.83    1.00   65.0
## behavior*            7 34    2.00  0.00    2.00    2.00  0.00    2.00    2.0
## behavior_daytime*    8 34    4.00  0.00    4.00    4.00  0.00    4.00    4.0
## first                9 34    0.68  0.47    1.00    0.71  0.00    0.00    1.0
## size_cm             10 23   11.85  1.71   11.61   11.73  1.42    8.71   16.4
## reaction            11 34    0.94  0.24    1.00    1.00  0.00    0.00    1.0
## responselatency1    12 32    3.38  1.52    3.00    3.23  1.04    1.20    8.7
## reaction2           13  2    1.00  0.00    1.00    1.00  0.00    1.00    1.0
## responselatency2    14  2    2.40  0.71    2.40    2.40  0.74    1.90    2.9
## pot_sleep           15  2    1.00  0.00    1.00    1.00  0.00    1.00    1.0
## comments*           16 34    1.32  1.07    1.00    1.00  0.00    1.00    5.0
##                    range  skew kurtosis   se
## date*               1.00  1.91     1.68 0.06
## temp                1.00  1.91     1.68 0.06
## location*           0.00   NaN      NaN 0.00
## time              187.00 -0.03    -0.78 8.40
## daytime*            0.00   NaN      NaN 0.00
## subject*           64.00  0.96     0.31 3.01
## behavior*           0.00   NaN      NaN 0.00
## behavior_daytime*   0.00   NaN      NaN 0.00
## first               1.00 -0.72    -1.52 0.08
## size_cm             7.69  0.69     0.68 0.36
## reaction            1.00 -3.59    11.19 0.04
## responselatency1    7.50  1.40     2.47 0.27
## reaction2           0.00   NaN      NaN 0.00
## responselatency2    1.00  0.00    -2.75 0.50
## pot_sleep           0.00   NaN      NaN 0.00
## comments*           4.00  2.86     6.53 0.18
```

```
describeBy(a$responselatency1, a$behavior_daytime)
```

```
##
##  Descriptive statistics by group
## group: active_day
##    vars  n mean   sd median trimmed  mad min max range skew kurtosis   se
## X1    1 12 1.58 0.47    1.5    1.55 0.59 1.1 2.4   1.3 0.48    -1.37 0.14
## ------------------------------------------------------------
## group: active_night
##    vars  n mean   sd median trimmed mad min max range skew kurtosis   se
## X1    1 13 1.55 0.33    1.5    1.52 0.3 1.2 2.3   1.1 0.72    -0.39 0.09
## ------------------------------------------------------------
## group: immobile_day
##    vars  n mean   sd median trimmed  mad min max range skew kurtosis   se
## X1    1 30 2.05 0.88    1.8    1.92 0.74 1.1 4.8   3.7 1.22     1.13 0.16
## ------------------------------------------------------------
```

```
## group: immobile_night
##    vars  n mean   sd median trimmed  mad min max range skew kurtosis   se
## X1    1 32 3.38 1.52      3    3.23 1.04 1.2 8.7   7.5  1.4     2.47 0.27
```

#### Preliminary tests

Testing effects of size and test order

```r
size <- glmmTMB(responselatency1 ~ size_cm + (1|subject), data = a, family = Gamma)
simres <- simulateResiduals(size)
plot(simres)
```

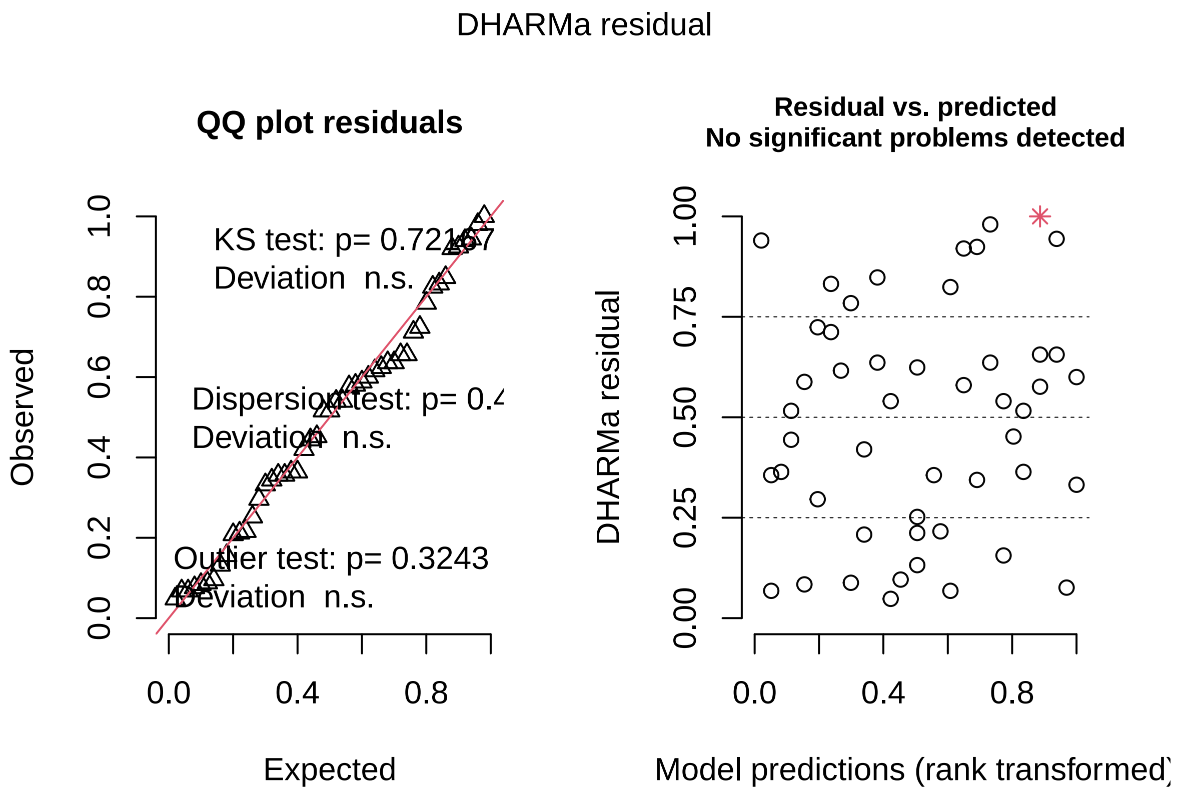


```r
Anova(size)
```

```
## Analysis of Deviance Table (Type II Wald chisquare tests)
##
## Response: responselatency1
##          Chisq Df Pr(>Chisq)
## size_cm 0.0936  1     0.7597
```

```r
summary(size)
```

```
##  Family: Gamma  ( inverse )
```

```
## Formula:          responselatency1 ~ size_cm + (1 | subject)
## Data: a
##
##      AIC      BIC   logLik deviance df.resid
##    161.2    168.8    -76.6    153.2       45
##
## Random effects:
##
## Conditional model:
##  Groups  Name        Variance Std.Dev.
##  subject (Intercept) 0.004576 0.06765
## Number of obs: 49, groups:  subject, 28
##
## Dispersion estimate for Gamma family (sigma^2): 0.175
##
## Conditional model:
##              Estimate Std. Error z value Pr(>|z|)
## (Intercept)  0.440890   0.200307   2.201   0.0277 *
## size_cm     -0.005202   0.017004  -0.306   0.7597
## ---
## Signif. codes:  0 '***' 0.001 '**' 0.01 '*' 0.05 '.' 0.1 ' ' 1
```

there is no effect of size on the response latency towards the first stimulus presentation

```r
order <- glmmTMB(responselatency1 ~ first + (1|subject), data = rcap, family = Gamma(link = "log"))
simres <- simulateResiduals(order)
plot(simres)
```

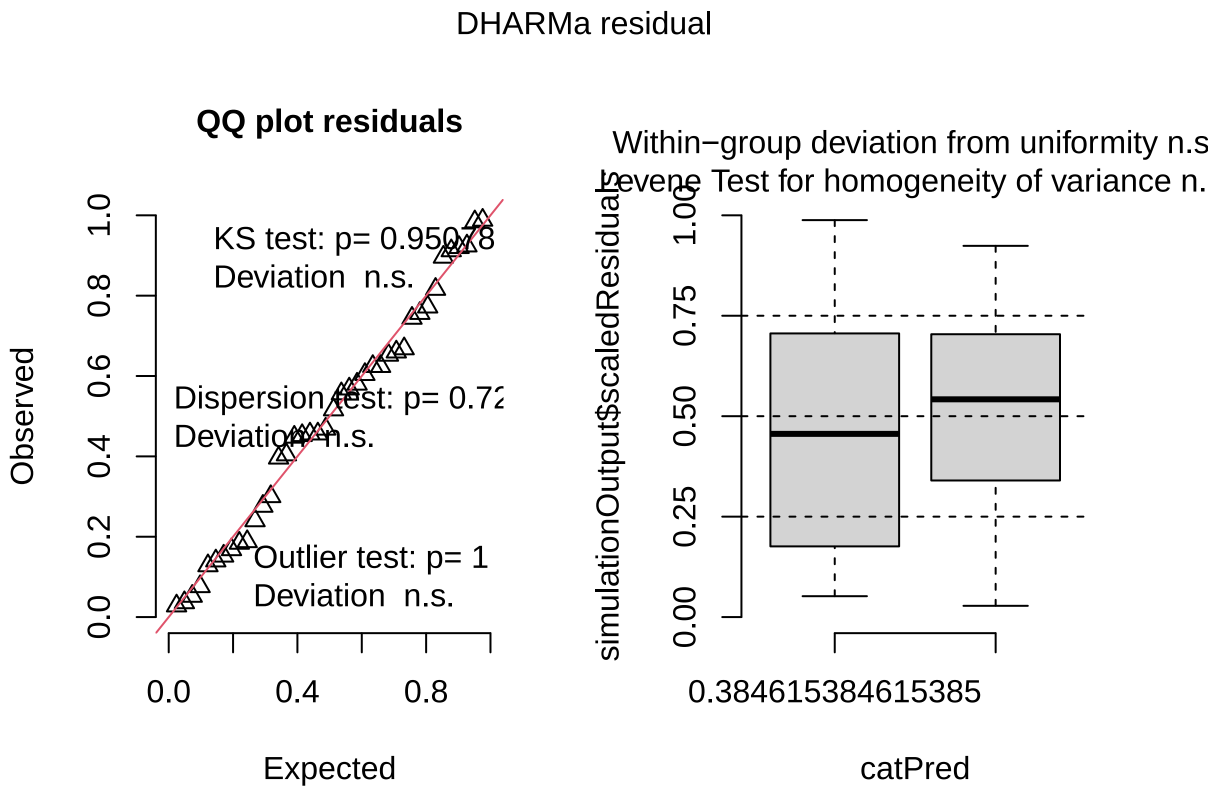


```r
Anova(order)
```

```
## Analysis of Deviance Table (Type II Wald chisquare tests)
##
## Response: responselatency1
##        Chisq Df Pr(>Chisq)
## first 1.497  1     0.2211
```

```r
summary(order)
```

```
##  Family: Gamma  ( log )
## Formula:          responselatency1 ~ first + (1 | subject)
## Data: rcap
##
##      AIC      BIC   logLik deviance df.resid
##    125.8    132.6    -58.9    117.8       36
##
## Random effects:
##
## Conditional model:
##  Groups  Name        Variance  Std.Dev.
##  subject (Intercept) 7.816e-10 2.796e-05
## Number of obs: 40, groups:  subject, 20
##
## Dispersion estimate for Gamma family (sigma^2): 0.171
```

```
##
## Conditional model:
##             Estimate Std. Error z value Pr(>|z|)
## (Intercept)   1.0944     0.1034  10.589   <2e-16 ***
## first        -0.1633     0.1334  -1.224    0.221
## ---
## Signif. codes:  0 '***' 0.001 '**' 0.01 '*' 0.05 '.' 0.1 ' ' 1
```

there is no effect of test order (night or day first) on the response latency towards the first stimulus presentation

### Main analyses

effect of daytime on response latency towards first stimulus presentation

```r
test <- glmmTMB(responselatency1 ~ daytime + (1|subject), data = a, family = Gamma(link="log"))
simres <- simulateResiduals(test)
plot(simres)
```

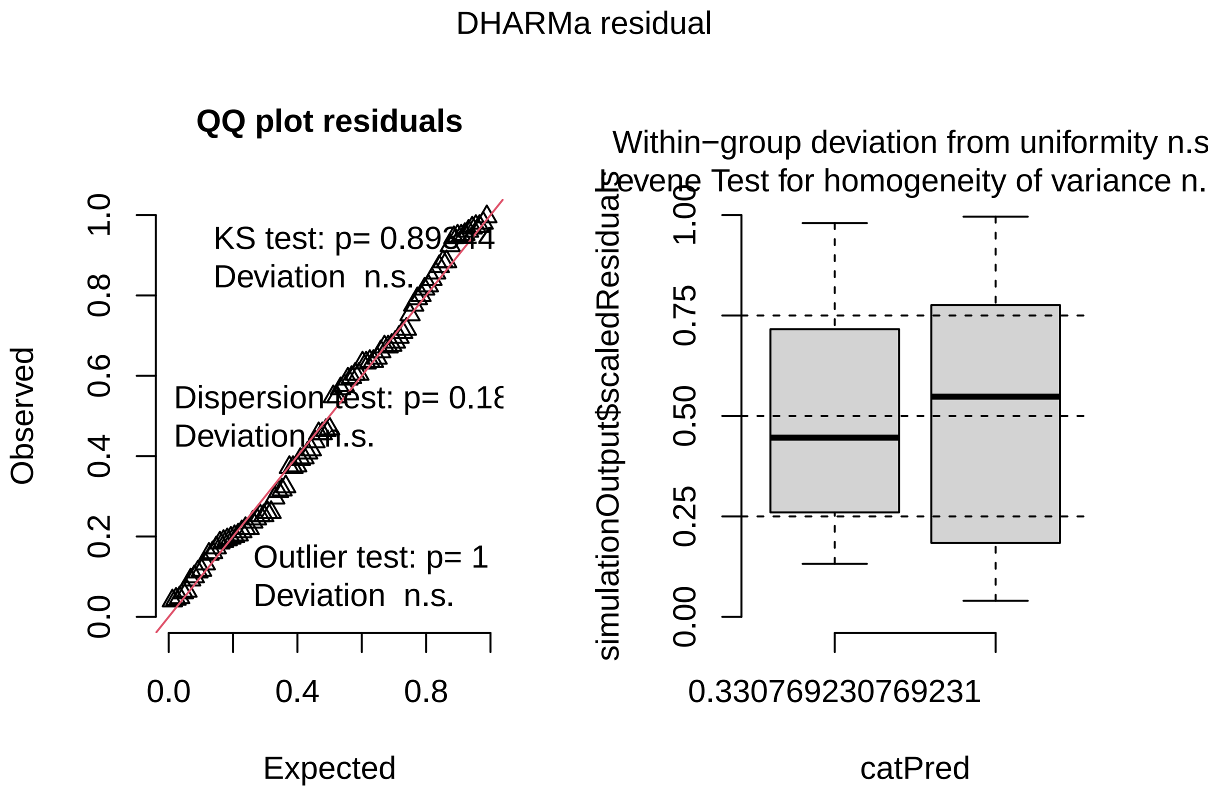


```r
summary(test)
```

```
##  Family: Gamma  ( log )
## Formula:          responselatency1 ~ daytime + (1 | subject)
## Data: a
```

```
##
##      AIC      BIC   logLik deviance df.resid
##    238.7    248.6   -115.3    230.7       83
##
## Random effects:
##
## Conditional model:
##  Groups  Name        Variance Std.Dev.
##  subject (Intercept) 0.0808   0.2842
## Number of obs: 87, groups:  subject, 61
##
## Dispersion estimate for Gamma family (sigma^2): 0.109
##
## Conditional model:
##              Estimate Std. Error z value Pr(>|z|)
## (Intercept)   0.59516    0.06807   8.743  < 2e-16 ***
## daytimenight  0.37504    0.07897   4.749 2.05e-06 ***
## ---
## Signif. codes:  0 '***' 0.001 '**' 0.01 '*' 0.05 '.' 0.1 ' ' 1
```

```r
Anova(test)
```

```
## Analysis of Deviance Table (Type II Wald chisquare tests)
##
## Response: responselatency1
##          Chisq Df Pr(>Chisq)
## daytime 22.551  1  2.046e-06 ***
## ---
## Signif. codes:  0 '***' 0.001 '**' 0.01 '*' 0.05 '.' 0.1 ' ' 1
```

there is a significant effect of daytime on the response latency

next we test the effect of behavior and daytime on the response latency

```r
test2 <- glmmTMB(responselatency1 ~ behavior_daytime + (1|subject), data = a, family = Gamma(link="log")
simres <- simulateResiduals(test2)
plot(simres)
```

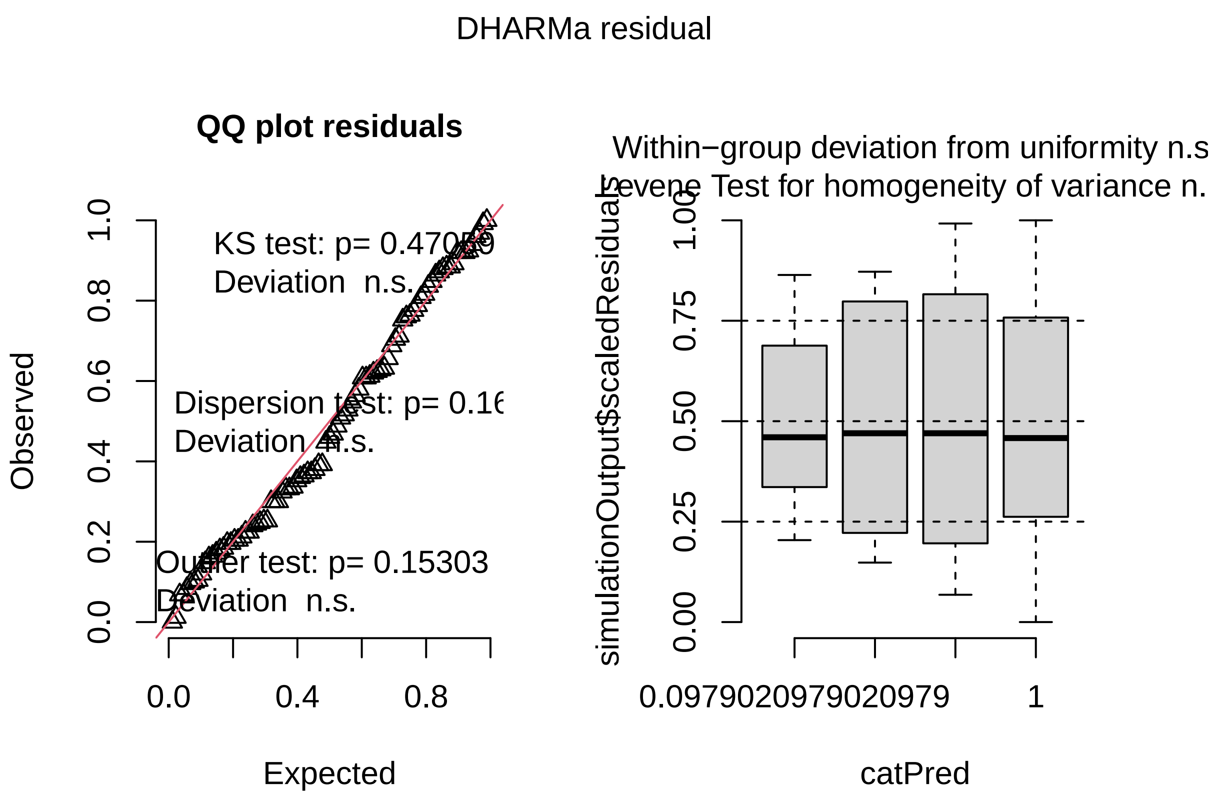


```r
summary(test2)
```

```
##  Family: Gamma  ( log )
## Formula:          responselatency1 ~ behavior_daytime + (1 | subject)
## Data: a
##
##      AIC      BIC   logLik deviance df.resid
##    213.2    228.0   -100.6    201.2       81
##
## Random effects:
##
## Conditional model:
##  Groups  Name        Variance Std.Dev.
##  subject (Intercept) 0.04034  0.2009
## Number of obs: 87, groups:  subject, 61
##
## Dispersion estimate for Gamma family (sigma^2): 0.0906
##
## Conditional model:
##                                 Estimate Std. Error z value Pr(>|z|)
## (Intercept)                      0.47400    0.10403   4.556 5.20e-06 ***
## behavior_daytimeactive_night    -0.02512    0.13896  -0.181   0.8566
## behavior_daytimeimmobile_day     0.20615    0.12383   1.665   0.0959 .
## behavior_daytimeimmobile_night   0.69557    0.12391   5.614 1.98e-08 ***
## ---
## Signif. codes:  0 '***' 0.001 '**' 0.01 '*' 0.05 '.' 0.1 ' ' 1
```

```r
Anova(test2)
```

```
## Analysis of Deviance Table (Type II Wald chisquare tests)
##
## Response: responselatency1
##                  Chisq Df Pr(>Chisq)
## behavior_daytime 62.22  3  1.971e-13 ***
## ---
## Signif. codes:  0 '***' 0.001 '**' 0.01 '*' 0.05 '.' 0.1 ' ' 1
```

```r
#posthoc test, comparisons
e <- emmeans(test2, ~ behavior_daytime, type="response")
pairs(e, adjust = "bonferroni")
```

```
##  contrast                      ratio     SE  df null z.ratio p.value
##  active_day / active_night     1.025 0.1425 Inf    1   0.181  1.0000
##  active_day / immobile_day     0.814 0.1008 Inf    1  -1.665  0.5757
##  active_day / immobile_night   0.499 0.0618 Inf    1  -5.614  <.0001
##  active_night / immobile_day   0.794 0.0950 Inf    1  -1.932  0.3198
##  active_night / immobile_night 0.486 0.0582 Inf    1  -6.023  <.0001
##  immobile_day / immobile_night 0.613 0.0497 Inf    1  -6.034  <.0001
##
## P value adjustment: bonferroni method for 6 tests
## Tests are performed on the log scale
```

with nonresponders scored as 10s

```r
#transform all NAs to 10s
nonres <- a
nonres["responselatency1"][is.na(nonres["responselatency1"])] <- 10

#model
testX2 <- glmmTMB(responselatency1 ~ behavior_daytime + (1|subject), data = nonres, family = Gamma(link
simres <- simulateResiduals(testX2)
plot(simres)
```

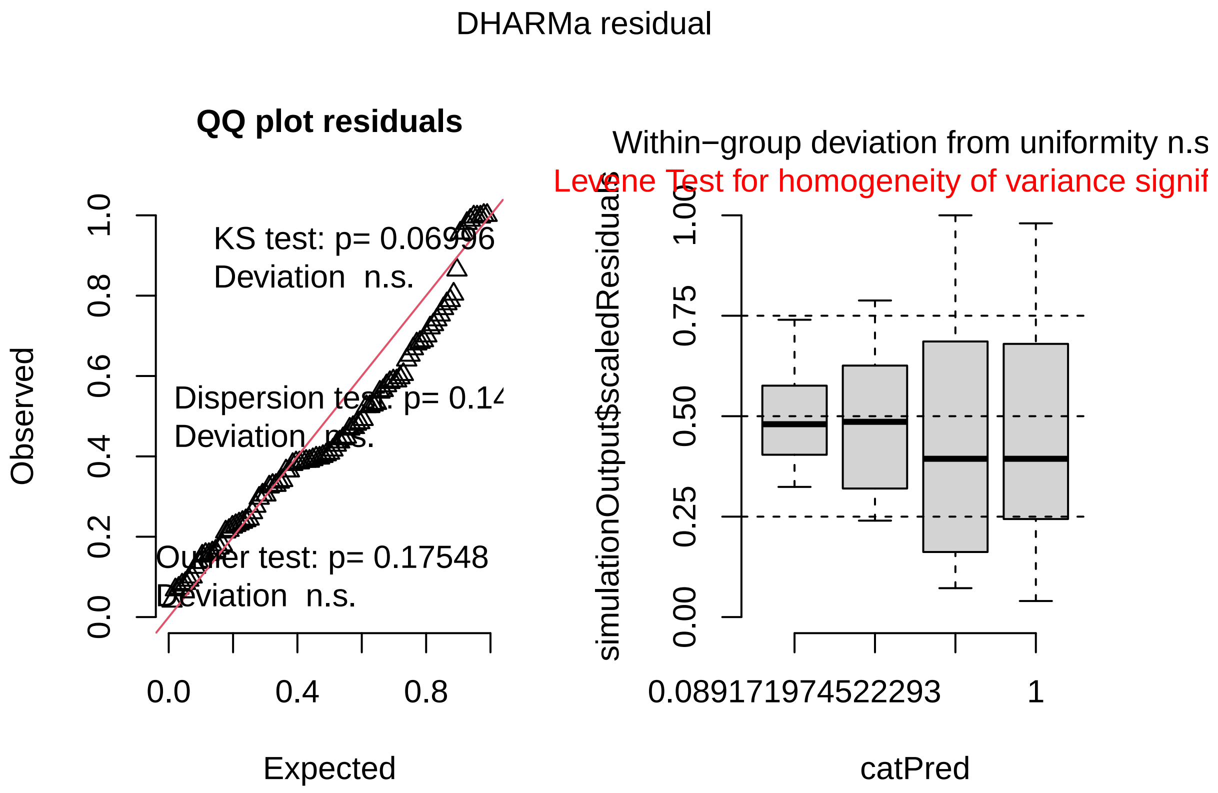


```r
summary(testX2)
```

```
##  Family: Gamma  ( log )
## Formula:          responselatency1 ~ behavior_daytime + (1 | subject)
## Data: nonres
##
##      AIC      BIC   logLik deviance df.resid
##    338.7    354.1   -163.4    326.7       89
##
## Random effects:
##
## Conditional model:
##  Groups  Name        Variance Std.Dev.
##  subject (Intercept) 0.1672   0.4089
## Number of obs: 95, groups:  subject, 67
##
## Dispersion estimate for Gamma family (sigma^2): 0.158
##
## Conditional model:
##                                 Estimate Std. Error z value Pr(>|z|)
## (Intercept)                      0.51731    0.16022   3.229  0.00124 **
## behavior_daytimeactive_night    -0.03718    0.20589  -0.181  0.85670
## behavior_daytimeimmobile_day     0.47971    0.19024   2.522  0.01168 *
## behavior_daytimeimmobile_night   0.82512    0.18612   4.433 9.28e-06 ***
## ---
## Signif. codes:  0 '***' 0.001 '**' 0.01 '*' 0.05 '.' 0.1 ' ' 1
```

```r
Anova(testX2)
```

```
## Analysis of Deviance Table (Type II Wald chisquare tests)
##
## Response: responselatency1
##                  Chisq Df Pr(>Chisq)
## behavior_daytime 32.008  3  5.213e-07 ***
## ---
## Signif. codes:  0 '***' 0.001 '**' 0.01 '*' 0.05 '.' 0.1 ' ' 1
```

```r
#posthoc test, comparisons
e <- emmeans(testX2, ~ behavior_daytime, type="response")
pairs(e, adjust = "bonferroni")
```

```
##  contrast                      ratio     SE  df null z.ratio p.value
##  active_day / active_night     1.038 0.2137 Inf    1   0.181  1.0000
##  active_day / immobile_day     0.619 0.1177 Inf    1  -2.522  0.0701
##  active_day / immobile_night   0.438 0.0816 Inf    1  -4.433  0.0001
##  active_night / immobile_day   0.596 0.1093 Inf    1  -2.820  0.0288
##  active_night / immobile_night 0.422 0.0765 Inf    1  -4.757  <.0001
##  immobile_day / immobile_night 0.708 0.0827 Inf    1  -2.958  0.0186
##
## P value adjustment: bonferroni method for 6 tests
## Tests are performed on the log scale
```

```r
testX3 <- glmmTMB(reaction ~ behavior_daytime, data = a, family = binomial)
simres <- simulateResiduals(testX3)
plot(simres)
```

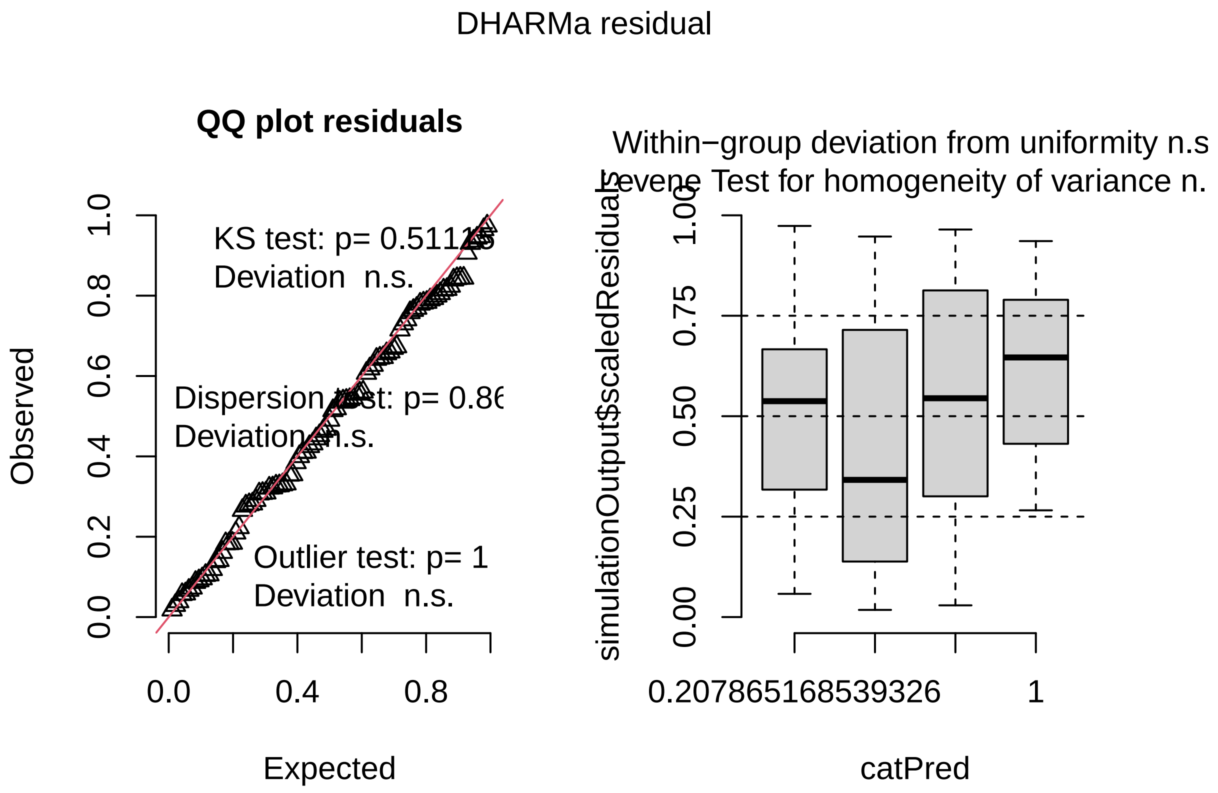


```r
summary(testX3)
```

```
##  Family: binomial  ( logit )
## Formula:          reaction ~ behavior_daytime
## Data: a
##
##      AIC      BIC   logLik deviance df.resid
##     55.7     65.9    -23.8     47.7       91
##
##
## Conditional model:
##                                Estimate Std. Error z value Pr(>|z|)
## (Intercept)                   2.219e+01  1.902e+04   0.001    0.999
## behavior_daytimeactive_night    9.112e+00  1.739e+06   0.000    1.000
## behavior_daytimeimmobile_day   -2.058e+01  1.902e+04  -0.001    0.999
## behavior_daytimeimmobile_night -1.942e+01  1.902e+04  -0.001    0.999
```

```r
Anova(testX3)
```

```
## Analysis of Deviance Table (Type II Wald chisquare tests)
##
## Response: reaction
##                   Chisq Df Pr(>Chisq)
## behavior_daytime 1.8501  3     0.6041
```

```r
#posthoc test, comparisons
e <- emmeans(testX3, ~ behavior_daytime, type="response")
pairs(e, adjust = "bonferroni")
```

```
##  contrast                         odds.ratio       SE  df null z.ratio p.value
##  active_day / active_night          0.00e+00 1.92e+02 Inf    1   0.000  1.0000
##  active_day / immobile_day          8.68e+08 1.65e+13 Inf    1   0.001  1.0000
##  active_day / immobile_night        2.71e+08 5.16e+12 Inf    1   0.001  1.0000
##  active_night / immobile_day        7.87e+12 1.37e+19 Inf    1   0.000  1.0000
##  active_night / immobile_night      2.46e+12 4.28e+18 Inf    1   0.000  1.0000
##  immobile_day / immobile_night      0.00e+00 0.00e+00 Inf    1  -1.360  1.0000
##
## P value adjustment: bonferroni method for 6 tests
## Tests are performed on the log odds ratio scale
```

testing the effect of behavior and daytime exclusively for recaptured/retested spiders

```r
test3 <- glmmTMB(responselatency1 ~ behavior_daytime + (1|subject), data = rcap, family = Gamma(link="lo
simres <- simulateResiduals(test3)
plot(simres)
```

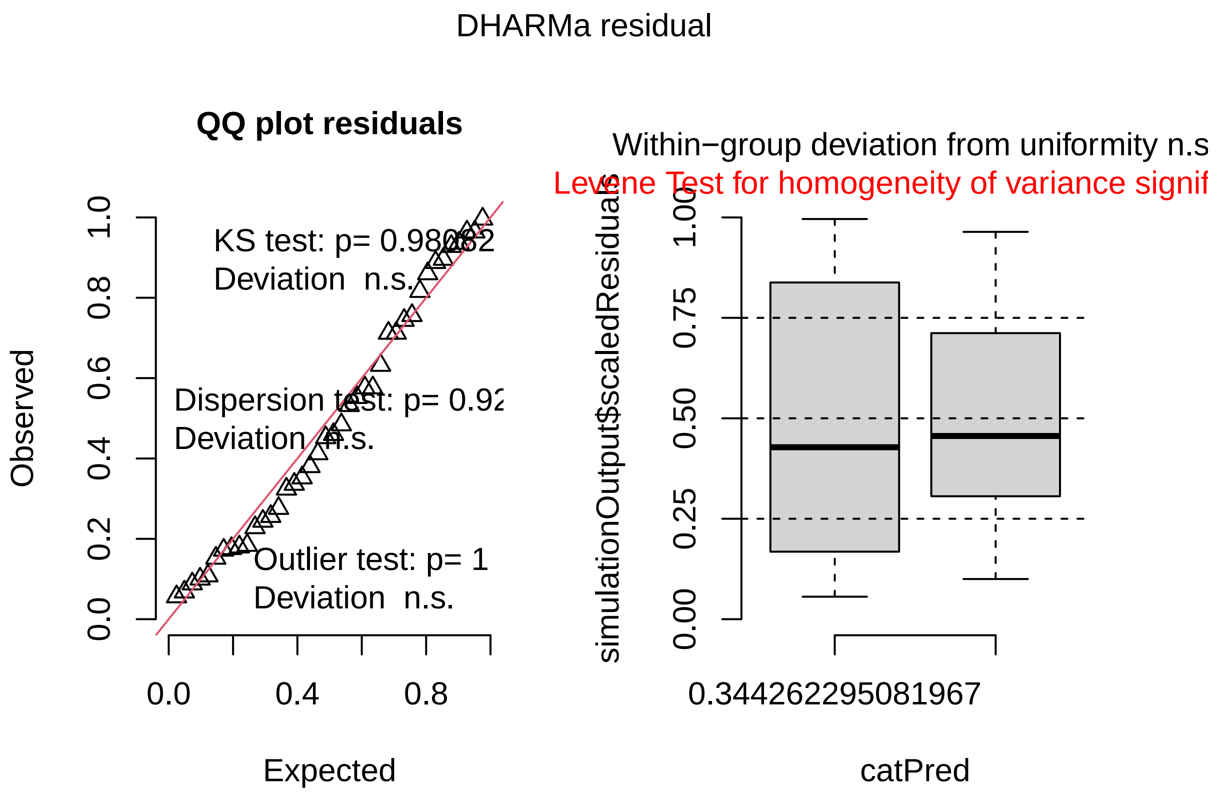


```r
summary(test3)
```

```
##  Family: Gamma  ( log )
```

```
## Formula:          responselatency1 ~ behavior_daytime + (1 | subject)
## Data: rcap
##
##      AIC      BIC   logLik deviance df.resid
##    112.8    119.6    -52.4    104.8       36
##
## Random effects:
##
## Conditional model:
##  Groups  Name        Variance Std.Dev.
##  subject (Intercept) 0.04747  0.2179
## Number of obs: 40, groups:  subject, 20
##
## Dispersion estimate for Gamma family (sigma^2): 0.0884
##
## Conditional model:
##                                  Estimate Std. Error z value Pr(>|z|)
## (Intercept)                       0.72728    0.08409   8.648  < 2e-16 ***
## behavior_daytimeimmobile_night  0.45230    0.09581   4.721 2.35e-06 ***
## ---
## Signif. codes:  0 '***' 0.001 '**' 0.01 '*' 0.05 '.' 0.1 ' ' 1
```

```r
Anova(test3)
```

```
## Analysis of Deviance Table (Type II Wald chisquare tests)
##
## Response: responselatency1
##                   Chisq Df Pr(>Chisq)
## behavior_daytime 22.285  1   2.35e-06 ***
## ---
## Signif. codes:  0 '***' 0.001 '**' 0.01 '*' 0.05 '.' 0.1 ' ' 1
```

the distribution is not perfect which is most likely due to the relatively small sample size of 20 individuals

### Plots

```r
#plot dB (total stimulus intensity is a product of dB and duration), with mean responselatencys for beh
db_plot <- ggplot(db, aes(responselatency1, stim))+
  geom_point()+
  geom_line()+
  geom_vline(xintercept = 1.58)+
  geom_vline(xintercept = 1.55)+
  geom_vline(xintercept = 2.05)+
  geom_vline(xintercept = 3.38)

db_plot
```

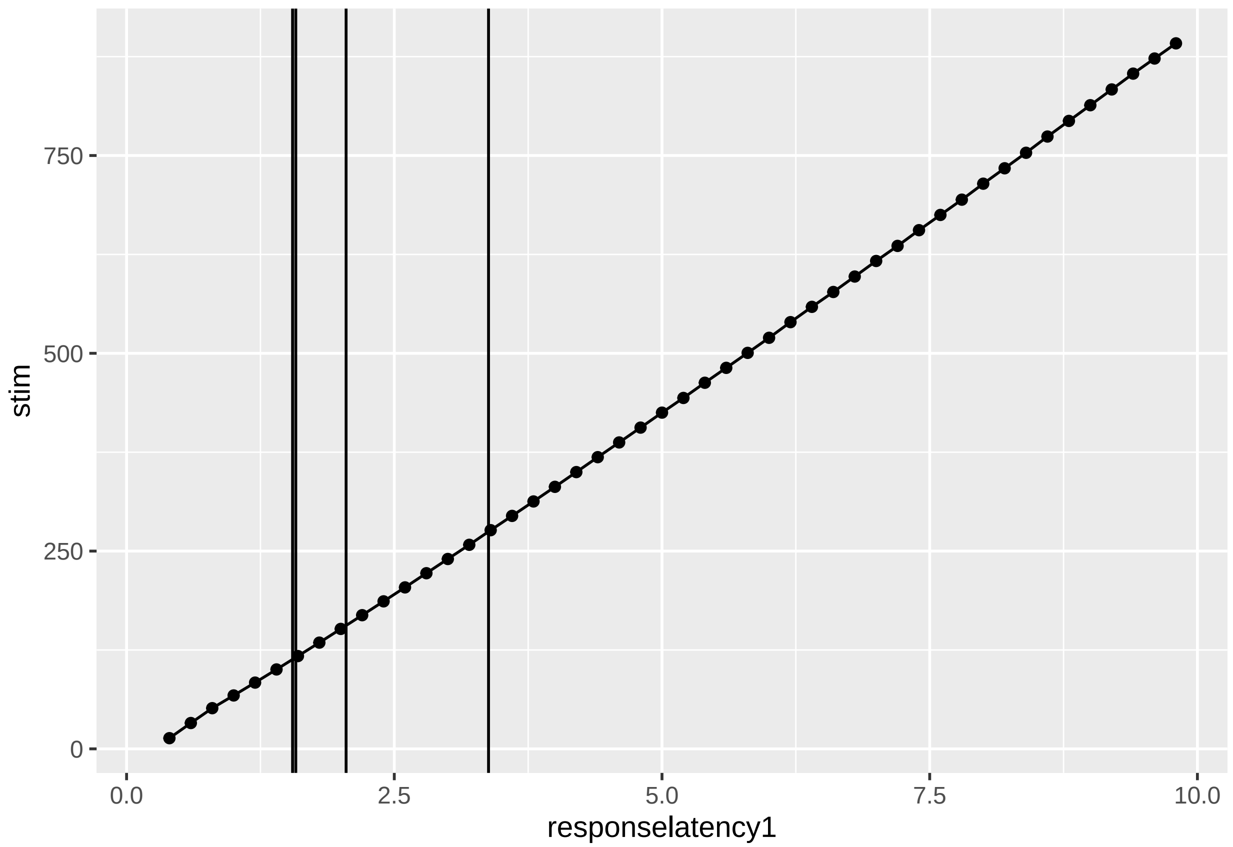


```
#ggsave("db_plot.svg", db_plot,  width = 5, height = 5, dpi = 300)
```

boxplots showing the response latency towards the first stimulus presentation for different behaviors and daytimes

```
arousal_boxplots <- ggplot(a, aes(behavior_daytime, responselatency1, fill = behavior_daytime))+
  geom_boxplot()+
  geom_jitter()+
  scale_fill_manual(values=cbbPalette)

arousal_boxplots
```

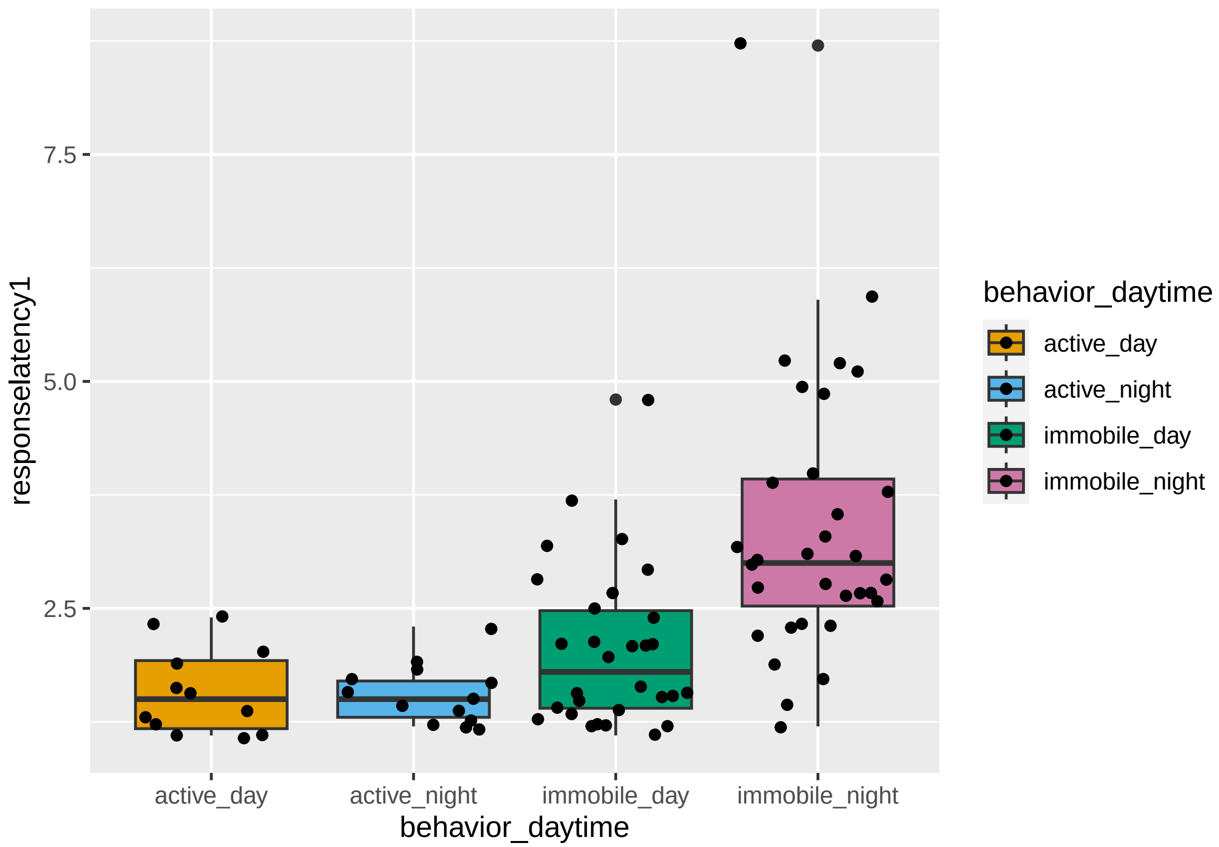


```
#ggsave("arousal_boxplots.svg", arousal_boxplots, width = 6, height = 5, dpi = 300)
```

with nonresponders

```
arousal_boxplots2 <- ggplot(nonres, aes(behavior_daytime, responselatency1, fill = behavior_daytime))+
  geom_boxplot()+
  geom_jitter()+
  scale_fill_manual(values=cbbPalette)

arousal_boxplots2
```

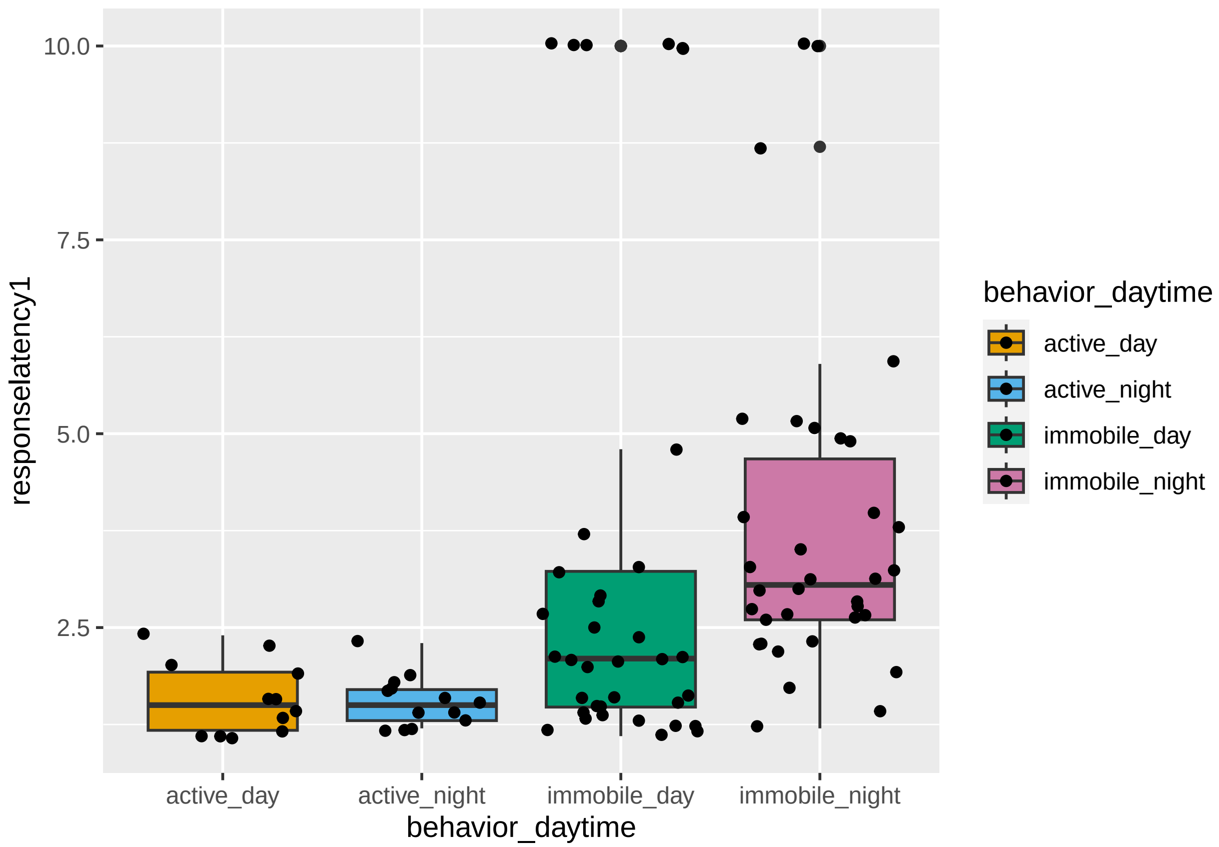


```
#ggsave("arousal_boxplots2.svg", arousal_boxplots2, width = 6, height = 5, dpi = 300)
```

boxplots showing the rectiontime towards the first stimulus presentation only for recaptured and retested spiders (both times during immobile states)

```
# with connecting lines between individuals
rcap_boxplot <- ggplot(rcap, aes(behavior_daytime, responselatency1, fill = daytime))+
  geom_boxplot(alpha = 0.5) +
   geom_line(aes(group = subject), size = .5) +
  geom_point(aes(group = subject), shape = 21, size = 3, color = 'white')+
  scale_colour_manual(values=cbbPalette)+
  scale_fill_manual(values=cbbPalette)


rcap_boxplot
```

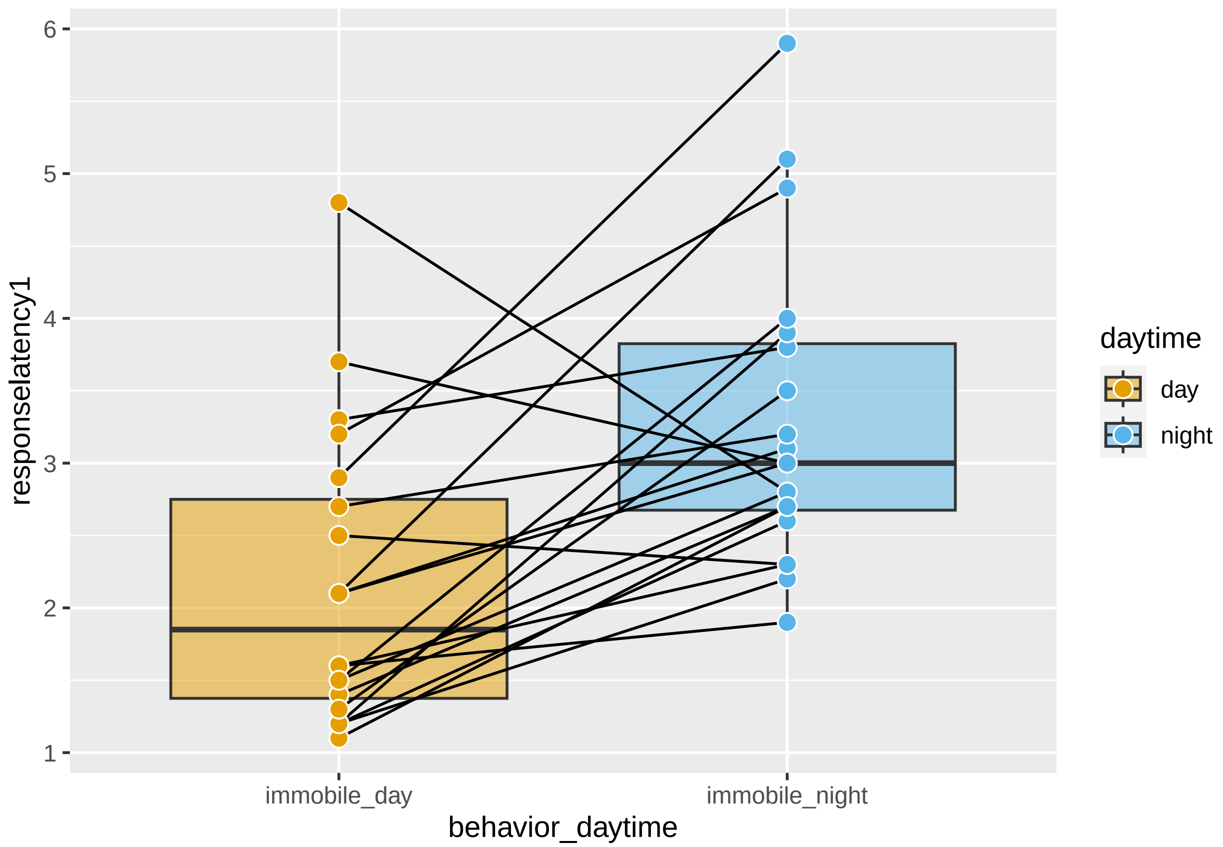


```
#ggsave("rcap_boxplot2.svg", rcap_boxplot , width = 5, height = 5, dpi = 300)
```

visualizing the frequency distribution of the response latency towards the first stimulus presentation based on behavior and daytime

```
density_plot <- ggplot(a, aes(responselatency1, behavior_daytime))+
  stat_density_ridges(quantile_lines = TRUE, quantiles = 2, scale = 2, alpha = .7)+
  xlim(0,9)

density_plot
```

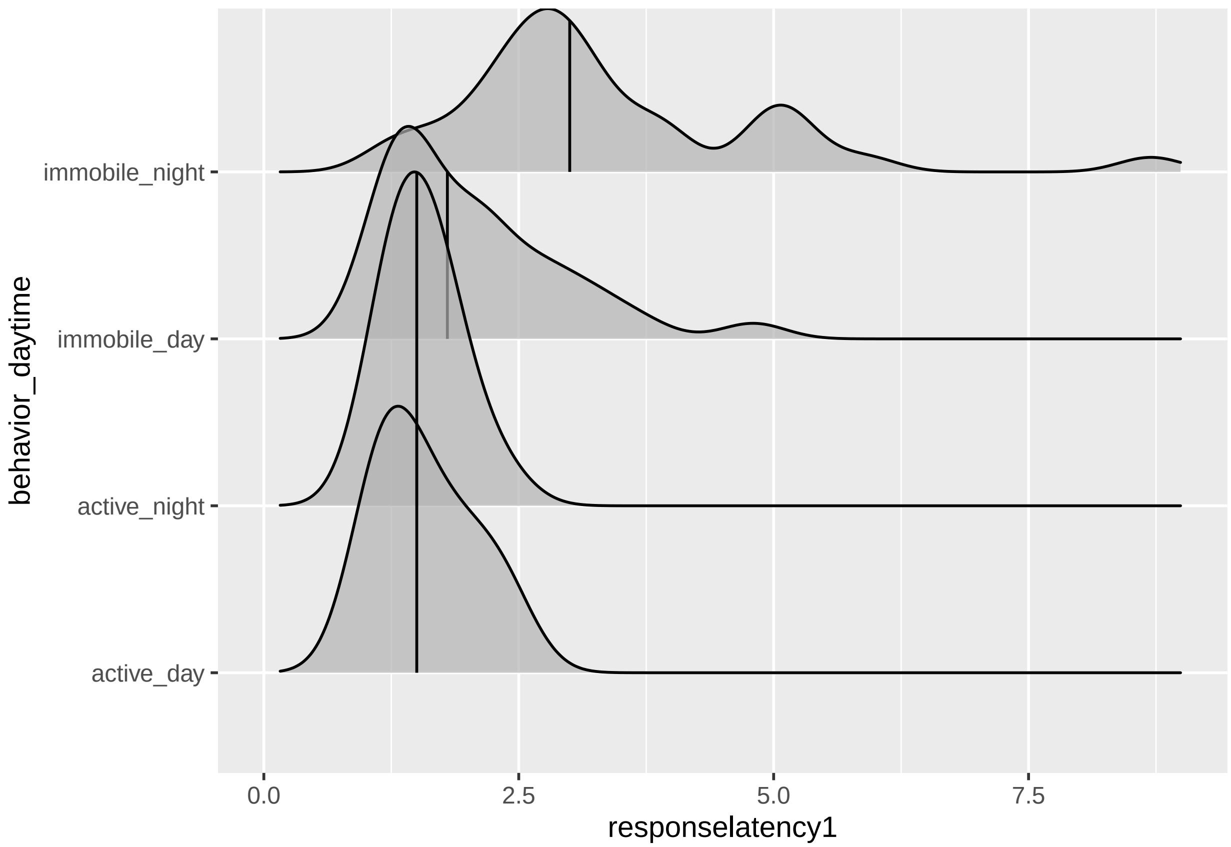


```
#ggsave("density_plot.svg", density_plot, width = 10, height = 5, dpi = 300)
```

with non responders included

```
density_plot2 <- ggplot(nonres, aes(responselatency1, behavior_daytime))+
  stat_density_ridges(quantile_lines = TRUE, quantiles = 2, scale = 2, alpha = .7)+
  xlim(0,11)

density_plot2
```

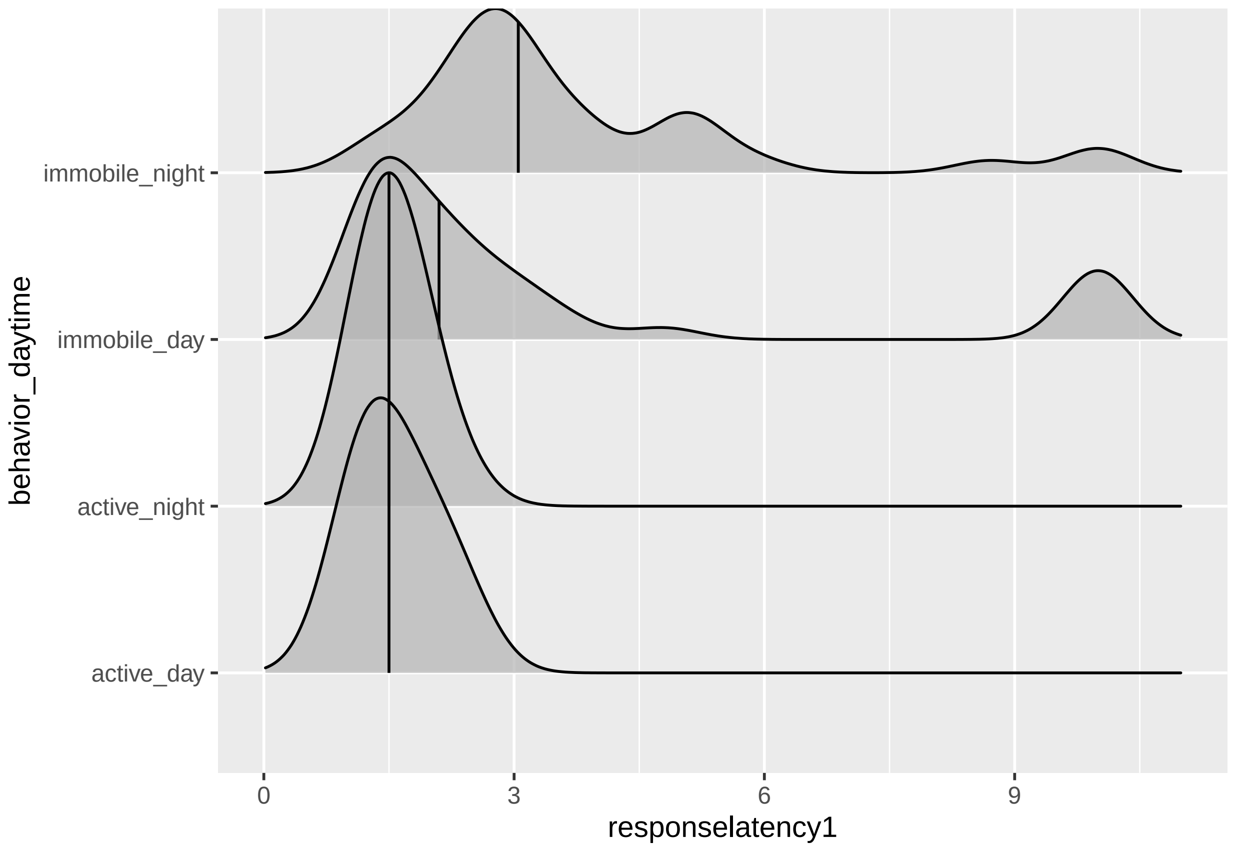


```r
ggsave("density_plot.svg", density_plot2, width = 10, height = 5, dpi = 300)
```

visualizing the response latency over the course of actual daytime for all tested spiders

```r
ggplot(a, aes(time, responselatency1))+
  geom_jitter()+
  geom_smooth(method="glm")
```

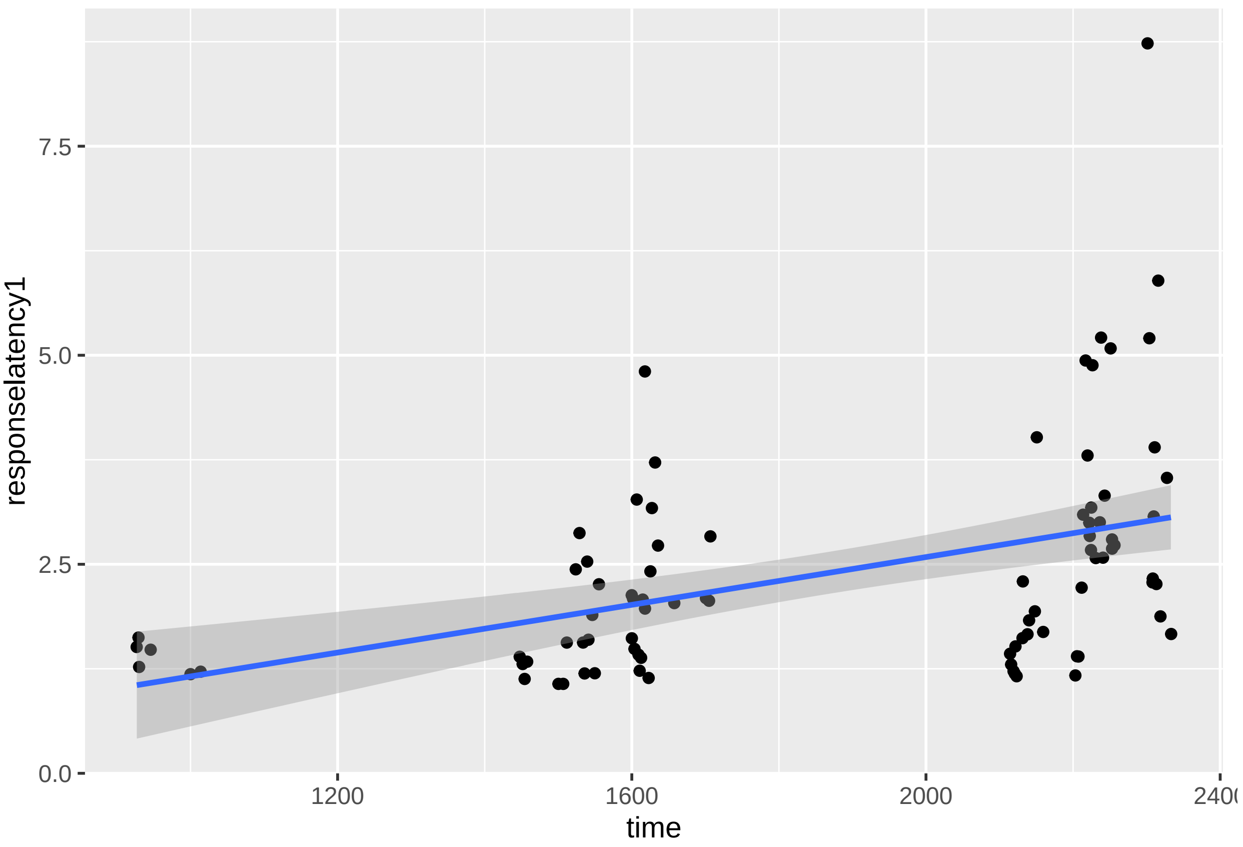
responselatency1
7.5
5.0
2.5
0.0
1200
1600
2000
2400
time
